# The global ocean size-spectrum from bacteria to whales

**DOI:** 10.1101/2021.04.03.438320

**Authors:** Ian A. Hatton, Ryan F. Heneghan, Yinon M. Bar-On, Eric D. Galbraith

**Affiliations:** Max Planck Institute for Mathematics in the Sciences, Leipzig 04103, Germany; Institut de Ciència i Tecnologia Ambientals (ICTA), Universitat Autonoma de Barcelona, Spain; School of Mathematical Sciences, Queensland University of Technology, Brisbane, Australia; Department of Plant and Environmental Sciences, Weizmann Institute of Science, 76100 Rehovot, Israel; Department of Earth and Planetary Sciences, McGill University, Montreal, Canada.

## Abstract

It has long been hypothesized that aquatic biomass is evenly distributed among logarithmic body mass size-classes. Although this community structure has been observed locally among plankton groups, its generality has never been formally tested across all marine life, nor have its impacts by humans been broadly assessed. Here, we bring together data at the global scale to test the hypothesis from bacteria to whales. We find that biomass within most order of magnitude size-classes is indeed remarkably constant, near 1 Gt wet weight (10^15^ grams), but that bacteria and whales are markedly above and below this value, respectively. Furthermore, human impacts have significantly truncated the upper one-third of the spectrum. Size-spectrum theory has yet to provide an explanation for what is possibly life’s largest scale regularity.

**One Sentence Summary:** Human activities have fundamentally altered one of life’s largest scale patterns; a global power law size distribution spanning bacteria to whales.

## Main Text

In 1972, Sheldon et al. published measurements of marine plankton abundance spanning six orders of magnitude in body mass (a factor of ∼100 in body length), collected at approximately 80 Atlantic and Pacific stations in a circum-navigation of the Americas. At each station the total biomass of all individuals was approximately evenly distributed across logarithmic size-classes (*1*). Based on these planktonic observations, they boldly hypothesized that “*to a first approximation, roughly equal concentrations of material occur at all particle sizes within the range from 1 µm to about 10^6^ µm, i.e., from bacteria to whales*”. This extraordinary hypothesis has never been formally tested across the astronomical range in body masses it encompasses, nor has its possible alteration by human impacts been examined at the whole-ocean scale.

Since Sheldon et al.’s seminal work, the distribution that results from aggregating individuals, regardless of species identity, into size bins has become known as the size-spectrum or Sheldon-spectrum, among other names (*2*). Although the ‘Sheldon hypothesis’ has been validated locally among plankton groups, often with striking similarity in reported exponents (near −1; Fig. S2) (*1*–*4*), it has not been examined globally, and has been limited to much smaller size ranges than the 23 orders of magnitude originally envisioned. Empirical size-spectra have typically averaged 6 orders of magnitude range in body mass, and have never exceeded 14 (Ref: (*4*); Table S3). Furthermore, the extent to which human activities may have impacted the whole-ocean size-spectrum has not been investigated.

Despite the proliferation of size-spectrum theory, (reviewed in (*2, 5–8*), there is still no accepted theoretical basis for why an even distribution of biomass should hold across all ocean life. Indeed, a great diversity of adaptive traits, from life-history and resource encounter strategies to mobility and sensory ability, depend on organisms “being the right size” (*7, 9–11*). This suggests that not all size-classes are created equal, and that certain sizes should be selectively advantaged or disadvantaged, which would suggest that the Sheldon hypothesis is mistaken or incomplete. Knowing the community structure of all marine life is /key to strengthening size-spectrum theory needed for a broader understanding of biosphere functioning and human impacts on the global ocean (*5, 8, 12*).

Here, we make use of advances in global ocean observation and recent meta-analyses to test Sheldon’s original hypothesis from bacteria to whales, at the global scale. We evaluate this hypothesis at a “pristine” time period before commercial-scale human harvesting of fish and marine mammals (pre-1800), as well as at present to evaluate possible alterations to the size-spectrum from human activity. Given the diversity of taxa and observational scales, we employed varied methods tailored to the different kinds of data available for each group. For example, phytoplankton are estimated globally using satellite images of surface chlorophyll, and converted to total depth-integrated biomass. Heterotrophic bacteria and all zooplankton groups, from single cells to large crustaceans, are estimated from >220,000 water samples, geographically distributed as shown in Fig. 1 A (black points), and interpolated over the whole ocean based on environmental correlates. The biomass of fish, which aggregate, migrate and can escape capture, is challenging to estimate from point samples, but are nonetheless intensively ‘sampled’ by commercial fisheries, so for fish biomass estimates we employ two independent global process models constrained by global catch data ((*13, 14*); Table S5). Finally, given that large marine mammals can individually range across whole ocean basins, we compile global population estimates for most marine mammal species (*n* = 82), and use body size allometry and geographic ranges to estimate the remainder (*n =* 44; Fig. S9).

**Fig 1.**
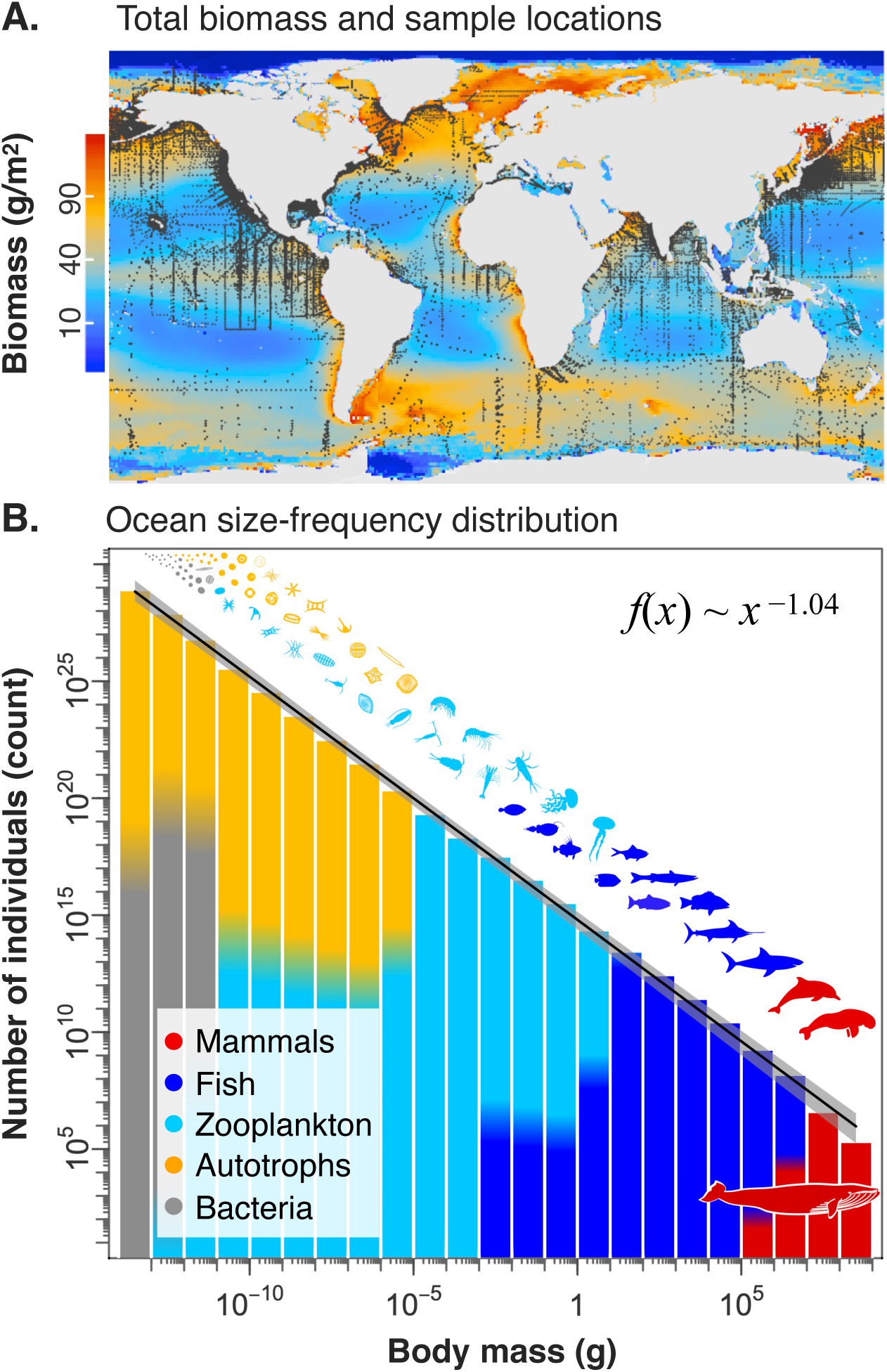
The global ocean size-spectrum. **A.** Biomass is estimated using different methods for different groups and summed over all groups in each 1° region of the ocean. The black points are *n* = 226,405 water sample locations for measurements of bacteria and zooplankton. **B**. The size-spectrum with total number of individuals in each order of magnitude size-class, estimated in the ocean’s epipelagic and continental shelves (upper ∼200 m), and giving an exponent −1.04 (95% C.I.: −1.05 to −1.02). The confidence band includes biomass uncertainty shown in Fig. 2 A, as well as uncertainty in the size distribution of each major group. Bin colors show the relative fraction of each group (no relation to y-axis, nor to the biomass in A).

These data and methods allow us to estimate the biomass of 12 major groups (aggregated into five groups in the Figures) over approximately 40,000 grid points of the global ocean. The total group biomasses are broadly concordant with prior global compilations of particular groups, as shown in Table S1 and S2 (*15–17*). We partition each group biomass into order of magnitude body mass size-classes based on published size distributions (Table S3), or if unknown, we partition biomass uniformly across the group size range (*18*). For each group and size-class we estimate a logarithmic 95% confidence interval, representing a multiplicative fold uncertainty, but we caution that uncertainty bounds are not strictly comparable across different methodologies. We also highlight conceptual uncertainty in delineating the appropriate bounds on the Sheldon hypothesis itself, which could include all ocean depths, estuary, benthic and sediment species, seabirds and polar bears. We address the various sources of uncertainty and their nuances in the SI (Table S4; (*18*)), and report here the most robust overall results.

We find that the estimated “pristine” global ocean size-spectrum is largely consistent with Sheldon’s original hypothesis (Fig. 1 B), particularly in the epipelagic (the upper, sunlit portion of the ocean). The observed slope, calculated from a regression fit to logarithmic size classes, is close to the long-hypothesized value of −1 (−1.04, 95% C.I.: −1.05 to −1.02), and exhibits remarkable regularity. This pattern indicates that biomass is generally not dominated by any best adapted size, as is evident when abundance is transformed to biomass (Fig. 2 A). However, our results show exceptions at the extremes: bacteria and whales diverge from the uniformity in biomass, which are more obvious when displayed on a linear scale (Fig. 2 B). Whereas all other groups sum to approximately 1 Gt wet weight (10^15^ grams) of biomass in each order of magnitude size bin, the size bins dominated by bacteria and whales are notably different. Although there is considerable uncertainty in our estimates (Fig. 2 A; (*18*)), these disparities are more pronounced and significant when we consider the size-spectrum over the full water column (cross-shading in Fig. 2), with bacteria dominating the biomass in the cold, dark ocean. Meso-pelagic nekton are also relatively abundant at depth (peak at 0.01 to 1 g) but these estimates are prone to large uncertainties (*15, 16, 18*). Sensitivity analysis shows that the overall size-spectrum slope is robust to both the choice of group biomass distributions, as well as the possible broad variations in biomass within our uncertainty bounds (slope 95% C.I. = ±0.043; Fig. 2 C i; Figs. S11 and S12) (*18*). Slope estimates are also robust to different fitting methods (Table S7), and are similar across global environmental gradients (Fig. S13).

**Fig 2.**
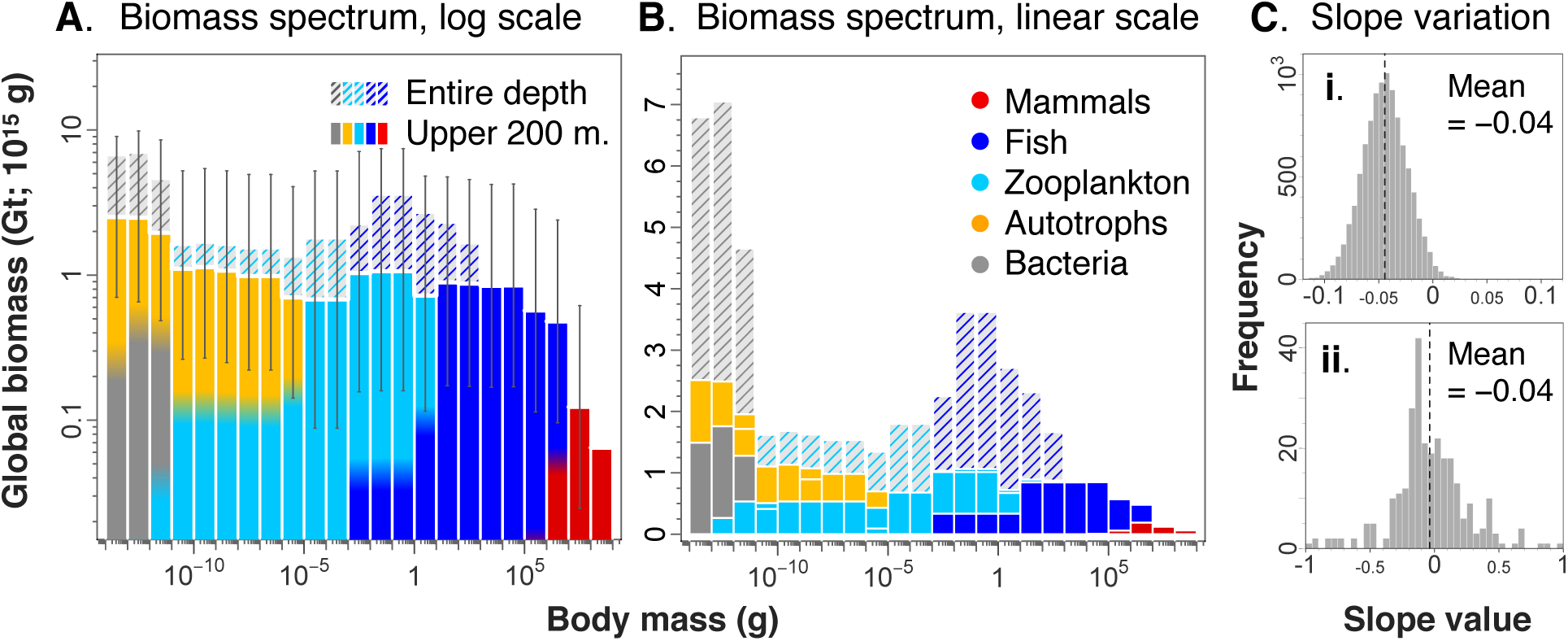
The ‘pristine’ ocean biomass spectrum. Total ocean biomass in each order of magnitude size-class is approximately 1 Gt (gigatonnes or petagrams = 10^15^ g), with exceptions at either extreme. Estimated historic biomass is shown in the upper 200 m of the ocean (colored), and extending to the seafloor (cross-shaded). Cross-shading is colored according to the group that dominates below the epipelagic, with bacteria dominating < 10^−11^ g, and meso-pelagic fish dominating size classes 10^−3^ to 10^3^ g). **A.** Global ocean biomass is shown on a logarithmic scale with logarithmic 95% confidence intervals on epipelagic biomass. Bin colors show the relative fraction of each group. **B.** Biomass estimates in A are shown on a linear scale to highlight differences of bacteria and whales from the overall trend. **C.** Frequency histograms of biomass spectrum slopes for **i**) resampled data incorporating our biomass uncertainty (*n* = 10,000 simulations); and **ii**) published slope values for *n* = 248 measured biomass spectra (from 41 separate studies; Ref: (*4*)).

Where might this law-like property of the “pristine” global ocean originate? An inverse relation between log size-class and frequency, as observed in Fig. 1, is a very simple pattern that describes things as different as cities (> 500,000 people; (*19*)) and asteroids (> 130 km diameter; (*20*)). In these cases, the size distribution is explained using a single abstract process such as random proportionate growth (*19*) or random fragmentation (*20*), that could apply to diverse phenomena. Size-spectrum theory, on the other hand, has tended towards greater biological realism, incorporating more factors, but at the expense of being able to identify a single dominant process responsible for the distribution. Existing theory has emphasized how incoming photosynthetic energy is partitioned according to the transfer of energy from one size-class (prey) to a larger size-class (predator), depending on individual rates of metabolism and consumption (reviewed in (*2, 5–8*). However, predator-prey mass ratios are not constant, and can vary up to eight orders of magnitude in any given size-class (Fig. S13) (*21–23*). Furthermore, recent work has shown that across this 23 order of magnitude size range, metabolism and consumption do not scale as ¾ with individual body size, as is often assumed, but closer to one (*10, 11, 24*). Other explanations for the size-spectrum have been based on the dynamics of how individuals grow across size-classes through ontogeny and reproduce smaller size-classes (*2, 5–8*). However, across groups there is enormous diversity in life-history, ranging from unicellular division, complex crustacean life cycles, fish egg spawning, and mammalian live birth (*25*). In sum, the size-spectrum appears more general and consistent than the individual level processes and allometries presumed to be its cause, leaving its origin a major unsolved problem.

This whole-ocean pattern is not immune to human impacts. Despite marine mammal and wild fish catches amounting to <3% of annual human food consumption (*18*), the cumulative impact that industrial fishing and whaling have had on the total biomass of these groups is striking. Fish >10 grams in size have been reduced in biomass by about 2 Gt (∼60% reduction; Fig. 3 A), and the largest size-classes have experienced a near 90% reduction in biomass since 1800 (Fig. 3 B). Using published impacts on major groups from high-emissions projected changes in climate (RCP 8.5), and assuming current fishing effort remains constant, we also estimate potential climatic impacts that could occur over the next century (Fig. 3 B). Our estimates suggest that fishing and whaling have already had a far greater impact among large size-classes than will climate change over the coming decades. The upper one-third of the biomass spectrum has been severely truncated, and the spectrum slope significantly altered (Fig. 3 C).

**Fig 3.**
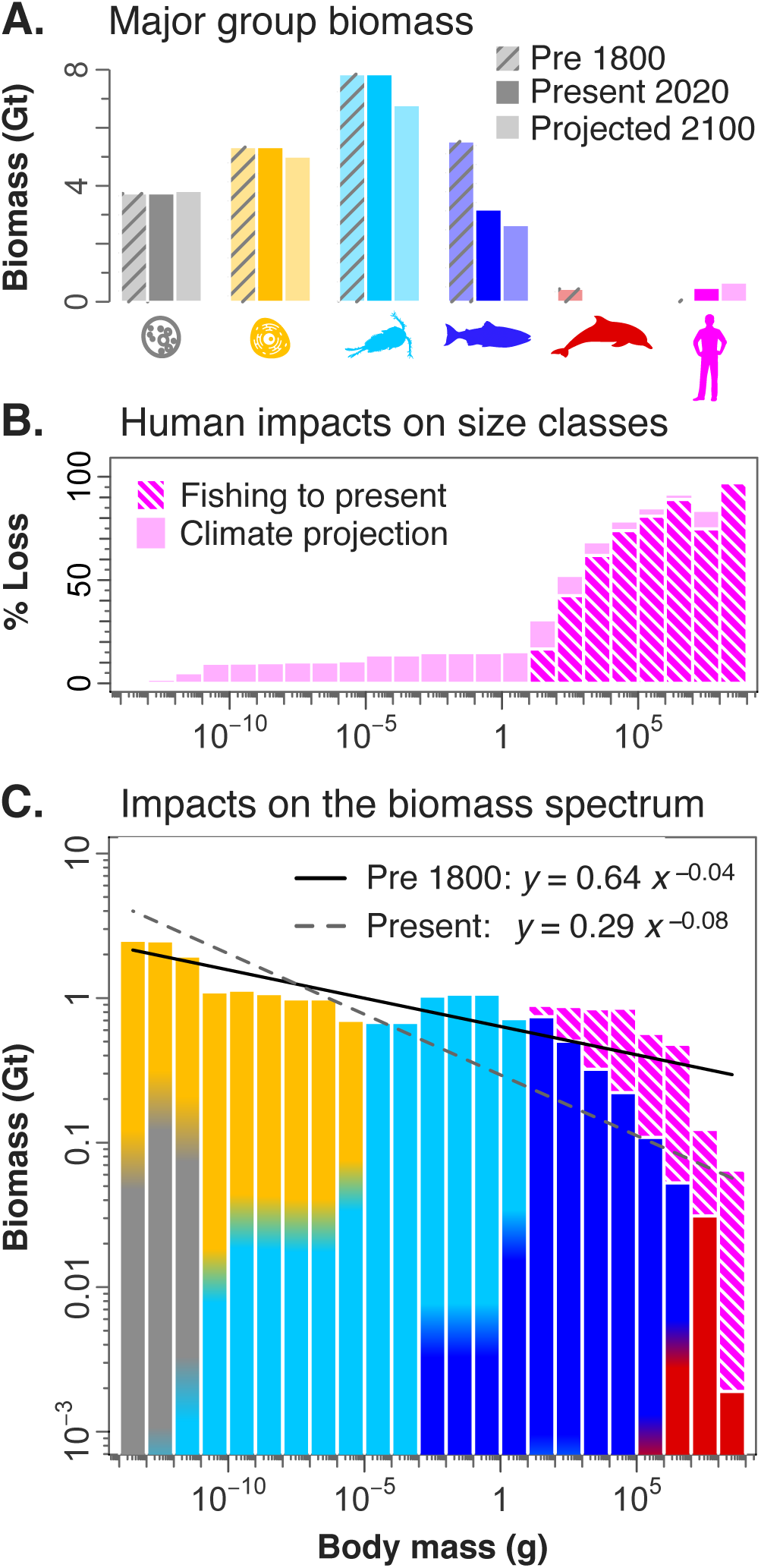
Human impacts on ocean biomass. **A**. Biomass in the ocean’s upper 200 m is compared across major groups at three time periods: before 1800 (hatched bars), current levels (solid bars) and projected to 2100 (light color), assuming similar fishing effort under extreme climate projections (RCP 8.5), based on published projected declines in major groups. **B**. Human impacts are estimated across size-classes, and show more extreme impacts above 10 grams. **C**. These impacts have altered the shape of the biomass spectrum, with the current day regression (2020; dashed grey line) having a significantly steeper slope than the pre-1800 regression (solid black line; “pristine”, as in Fig. 2).

Human biomass is now among the largest of any single species at approx. 0.4 Gt, and apparently surpassing Antarctic krill (*26*), long thought to be the most abundant naturally occurring animal species. This is, however, surprisingly small relative to the marine biomass lost from the largest size classes through human activity (∼2.7 Gt; Fig. 3 A). Since biomass is tightly coupled to production (*27*), the productivity of the largest size classes have been critically reduced, such that human catch consumption is on the order of 0.01 TW (10^12^ watts), compared to the roughly 1 TW energy production that would have otherwise been supported in a pristine system (*18*), and thus representing a near two order of magnitude loss in production efficiency among the size classes that humans value most.

Our results provide evidence that the “pristine” size-spectrum regularities previously observed at the local scale are largely preserved at the global scale, demanding renewed effort to uncover the processes underlying them. At the same time, our analysis has quantified a major impact of humanity on the distribution of biomass across size-ranges, highlighting the degree to which human activities in the Anthropocene have altered life at the global-scale.

## Acknowledgments

We thank A. Richardson, J. Everett, R. Milo, J. Guiet and A. Zadorin for early discussions, and J. Grady, M. Barbier and the BIGSEA Lab for feedback on an earlier draft. We are grateful for funding from the Alexander von Humboldt Foundation to I.H., the Spanish Ministry of Science, Innovation and Universities through the Acciones de Programación Conjunta Internacional to R.H. (PCIN-2017-115), and the European Research Council (ERC) under the European Union’s Horizon 2020 research and innovation programme to E.G. (grant agreement No. 682602).

The authors declare no competing financial interests.

## Supplementary Information

### Overview

To construct the global ocean size-spectrum, we drew on a diverse assemblage of complementary data, methods and assumptions. Here, we outline our data and methods for each group and our procedures used to reach the estimates of size-specific abundance, biomass and uncertainty values reported in the paper. We also detail the approach used to estimate how humans have altered the size-spectrum through animal hunting and climate change. Finally, we outline our statistical approach, including more general considerations of our treatment of uncertainty and fitting methods, as well as theoretical considerations, including the most common assumptions in size-spectrum theory and their foremost limitations. In addition, we provide all data and referenced sources, such that our analysis can be reproduced and updated as new data becomes available (https://github.com/ryanheneghan/sheldon_revisited.git).

There are several ways that a global size-spectrum analysis could be undertaken, from the coarse use of published global biomass estimates aggregated by major group and size class, to a very disaggregated assembly of existing observational data, which could include many tens of thousands of species-specific surveys and mass-specific size-spectra for different taxa and regions. The coarsely-aggregated extreme is shown in Table 1. In contrast, a highly disaggregated approach would require a vast effort, and would be plagued by difficulties regarding spatial and temporal bias. For this paper we elected to focus on a middle path between the two extremes, making use many kinds of pre-existing data, while employing a number of procedures of intermediate sophistication. Our methods are tailored to the data existing for each major taxonomic group, and are designed to maximize the utility of ancillary information.

Our methods include statistical and process models constrained by large existing surveys or satellite imagery, ensembles of dynamical models developed by independent research groups, and species specific survey data over their global geographic range. Our resulting global biomass estimates are all within a factor of two of previously published global biomass estimates (*15–17, 28, 29*) (Table S1). Abundance and biomass estimates for more resolved taxonomic groupings are shown in Table S2 and the geographic distribution of our biomass estimates for each major group is shown in Fig. S1. Finally, greater resolution and summary tables are available as part of the output of the code linked to in the first paragraph.

**Table S1.**
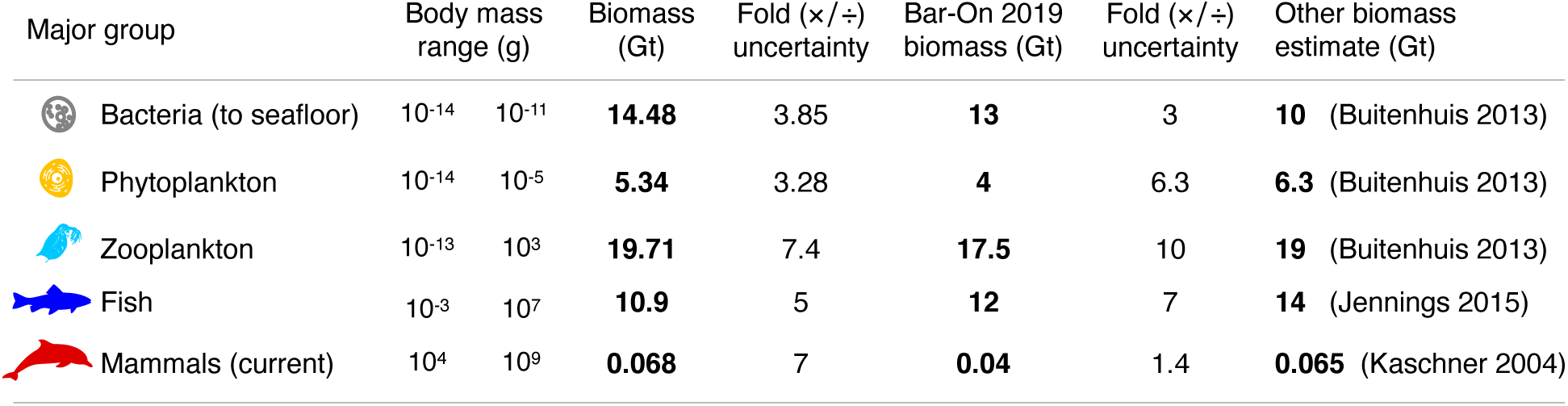
General summary of ocean biomass. Body mass range and total ocean biomass (wet weight; Gt ≡ 10^15^ g) are summarized across major taxonomic groups. Fold uncertainty is a multiplicative factor of the mean (×/÷) that expresses our confidence in the data akin to a 95% confidence interval (see text). These estimates are compared to values in the literature, particularly Bar-On et al. 2019 (*15–17, 28, 29*).

| Major group | Body mass range (g) | Biomass (Gt) | Fold ($\times/\div$) uncertainty | Bar-On 2019 biomass (Gt) | Fold ($\times/\div$) uncertainty | Other biomass estimate (Gt) |
| --- | --- | --- | --- | --- | --- | --- |
| Bacteria (to seafloor) | $10^{-14}$ $10^{-11}$ | **14.48** | 3.85 | **13** | 3 | **10** (Buitenhuis 2013) |
| Phytoplankton | $10^{-14}$ $10^{-5}$ | **5.34** | 3.28 | **4** | 6.3 | **6.3** (Buitenhuis 2013) |
| Zooplankton | $10^{-13}$ $10^3$ | **19.71** | 7.4 | **17.5** | 10 | **19** (Buitenhuis 2013) |
| Fish | $10^{-3}$ $10^7$ | **10.9** | 5 | **12** | 7 | **14** (Jennings 2015) |
| Mammals (current) | $10^4$ $10^9$ | **0.068** | 7 | **0.04** | 1.4 | **0.065** (Kaschner 2004) |

**Table S2.**
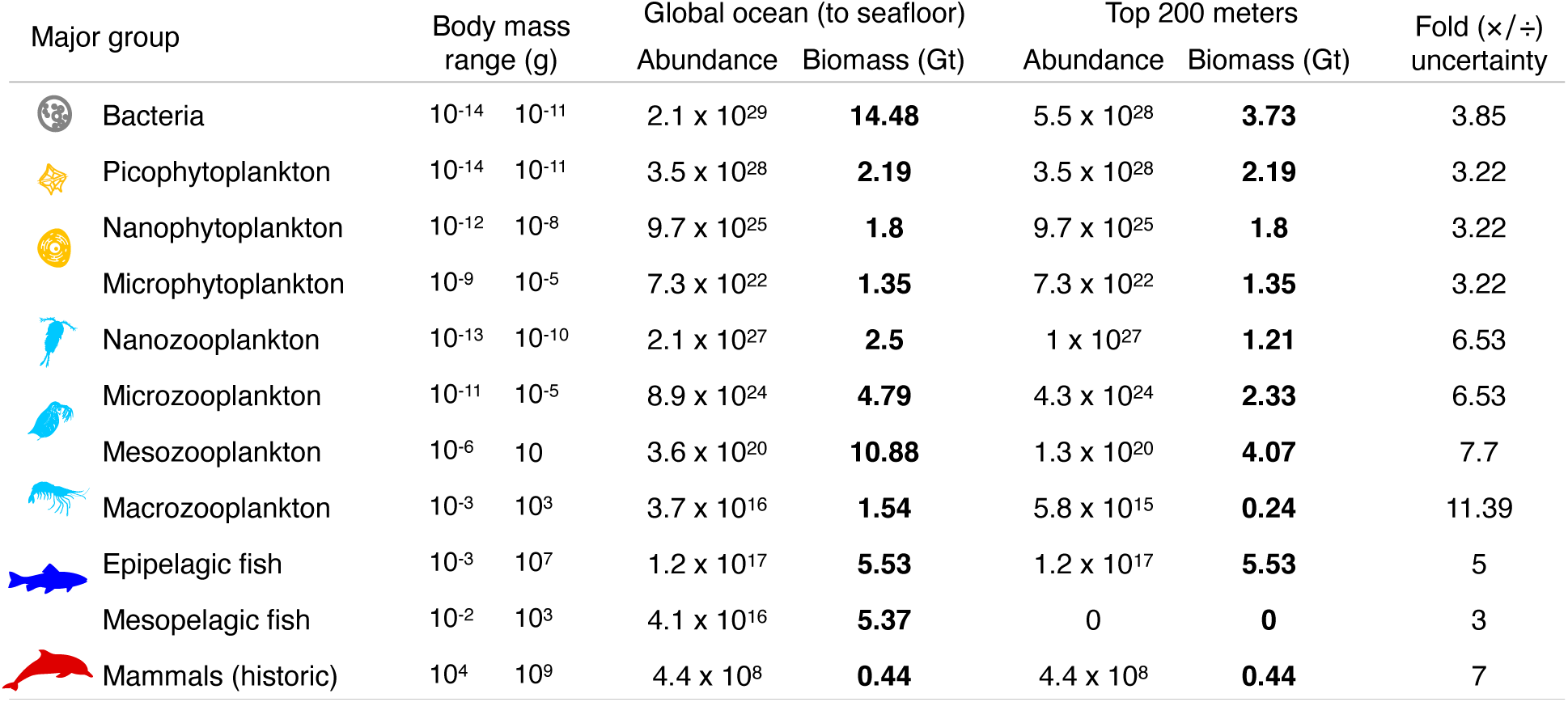
Summary of ocean biomass among major groups. Ocean abundance and biomass (wet weight) are compared across major groups at a more resolved level than Table S1 (see Table S1 caption), and include values for the top 200 meters and extended to the sea floor. Mammals and fish are estimated at historic levels.

| Major group | Body mass range (g) | | Global ocean (to seafloor) | | Top 200 meters | | Fold ($\times/\div$) uncertainty |
| --- | --- | --- | --- | --- | --- | --- | --- |
|  |  |  | Abundance | Biomass (Gt) | Abundance | Biomass (Gt) |  |
| Bacteria | 10<sup>-14</sup> | 10<sup>-11</sup> | 2.1 x 10<sup>29</sup> | **14.48** | 5.5 x 10<sup>28</sup> | **3.73** | 3.85 |
| Picophytoplankton | 10<sup>-14</sup> | 10<sup>-11</sup> | 3.5 x 10<sup>28</sup> | **2.19** | 3.5 x 10<sup>28</sup> | **2.19** | 3.22 |
| Nanophytoplankton | 10<sup>-12</sup> | 10<sup>-8</sup> | 9.7 x 10<sup>25</sup> | **1.8** | 9.7 x 10<sup>25</sup> | **1.8** | 3.22 |
| Microphytoplankton | 10<sup>-9</sup> | 10<sup>-5</sup> | 7.3 x 10<sup>22</sup> | **1.35** | 7.3 x 10<sup>22</sup> | **1.35** | 3.22 |
| Nanozooplankton | 10<sup>-13</sup> | 10<sup>-10</sup> | 2.1 x 10<sup>27</sup> | **2.5** | 1 x 10<sup>27</sup> | **1.21** | 6.53 |
| Microzooplankton | 10<sup>-11</sup> | 10<sup>-5</sup> | 8.9 x 10<sup>24</sup> | **4.79** | 4.3 x 10<sup>24</sup> | **2.33** | 6.53 |
| Mesozooplankton | 10<sup>-6</sup> | 10 | 3.6 x 10<sup>20</sup> | **10.88** | 1.3 x 10<sup>20</sup> | **4.07** | 7.7 |
| Macrozooplankton | 10<sup>-3</sup> | 10<sup>3</sup> | 3.7 x 10<sup>16</sup> | **1.54** | 5.8 x 10<sup>15</sup> | **0.24** | 11.39 |
| Epipelagic fish | 10<sup>-3</sup> | 10<sup>7</sup> | 1.2 x 10<sup>17</sup> | **5.53** | 1.2 x 10<sup>17</sup> | **5.53** | 5 |
| Mesopelagic fish | 10<sup>-2</sup> | 10<sup>3</sup> | 4.1 x 10<sup>16</sup> | **5.37** | 0 | **0** | 3 |
| Mammals (historic) | 10<sup>4</sup> | 10<sup>9</sup> | 4.4 x 10<sup>8</sup> | **0.44** | 4.4 x 10<sup>8</sup> | **0.44** | 7 |

**Fig. S1.**
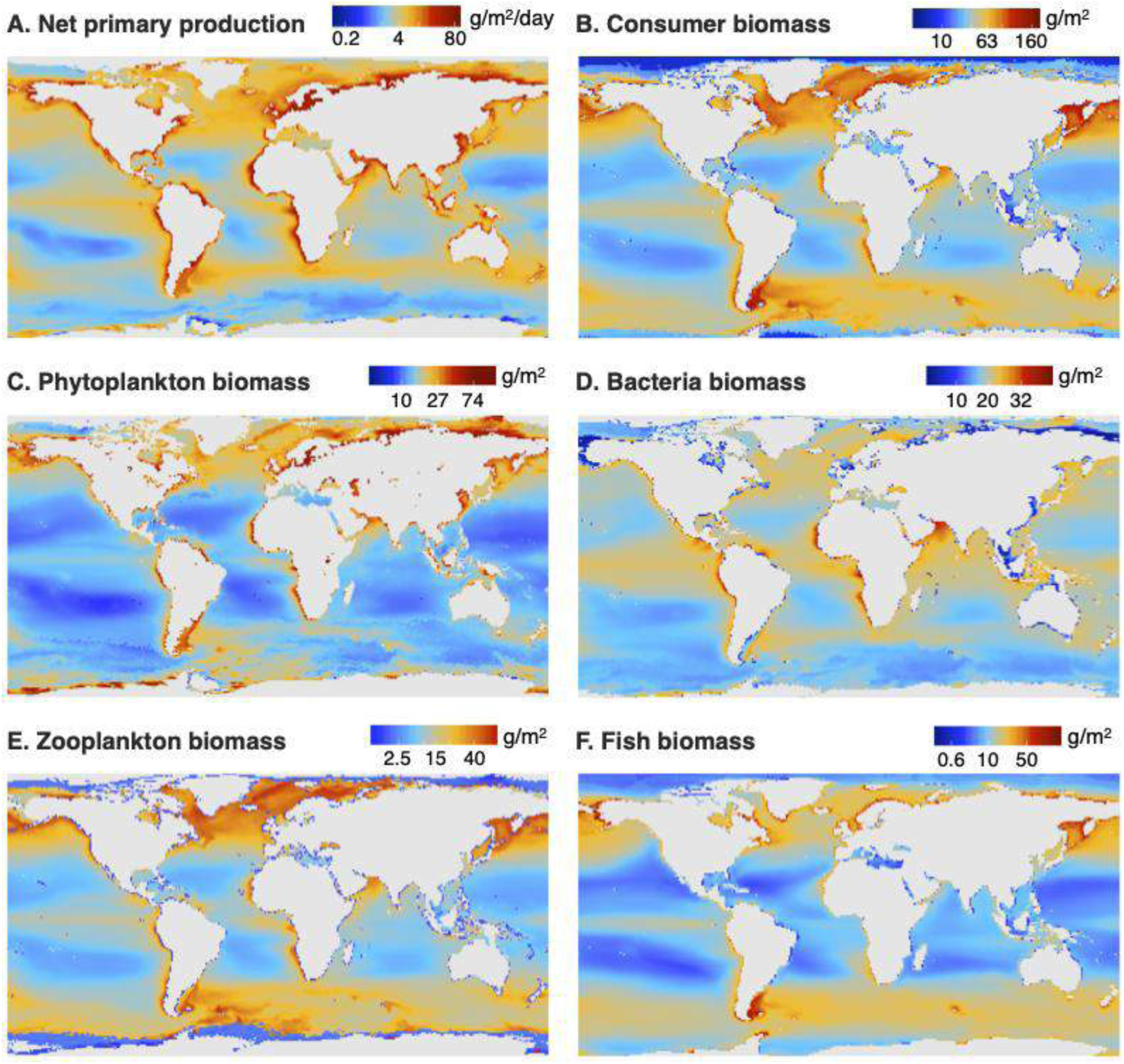
Geographic distribution of biomass by major taxonomic group. All biomass is for the top 200 m of the global ocean in g/m^2^. **A.** Net primary production (NPP) on which the vast majority of all other biomass is based. **B**. All consumer biomass is the sum of biomass in D to F that depends on the flux in A. **C.** Phytoplankton biomass is the standing stock that is largely responsible for NPP in A. **D-F.** Biomass of (D) bacteria, (E) zooplankton and (F) fish. Note the color scale of each major group is not the same, and was instead chosen to better distinguish the geographic distribution of biomass within a group, rather than facilitate cross-group comparison.

In order to study the size-spectrum over the entire ocean, we necessarily employed a number of assumptions to delimit the scope of our study, on which our results are strongly dependent. One major assumption is what constitutes “ocean” life. Like Sheldon and the many size-spectra studies that came after, we take oceanic life to refer largely to the pelagic environment (organisms living suspended in the salt water column of the ocean)(*1, 4, 30*). We therefore did not consider benthic organisms explicitly or in great depth, though aggregate measures of organisms that live on the seafloor are included in estimates of “non-mesopelagic fish” from the FishErIes Size and functional Type model (FEISTY; (*14*)) and the BiOeconomic mArine Trophic Size-spectrum model (BOATS; (*13*)). We also elected not to include brackish waters, mangroves or seabirds. The exclusion of benthic and sediment organisms has the largest impact on bacteria biomass estimates, given their extremely large estimated biomass in sediments (*31*).

We also did not explicitly consider several taxonomic groups including the following (global biomass estimates are from (*15, 16*): viruses (∼0.3 Gt); fungi (∼2.6 Gt); seagrasses (∼2 Gt), and corals (∼0.5 Gt). Our methods and estimates include groups such as molluscs including squid (0.5 Gt) and nematodes (0.1 Gt) that we did not separately or explicitly consider. We do not distinguish bacteria from archaea (3 Gt), and refer to these two groups as “bacteria”.

We show estimates of biomass over the top 200 meters of the ocean as well as the entire water column down to the sea floor to provide perspective on the vertical distribution of marine life. The top 200 meters (epipelagic) is where the vast majority of life’s diversity resides and where all major groups commonly exist, as well as where data have the best coverage. Extending our analysis from the top 200 meters to the ocean depths has the greatest effect on estimates of bacteria biomass (Table S2). Apart from phytoplankton and mammals, which only reside in surface waters, other groups tend to double their biomass going from the epipelagic to the seafloor (Table S2).

Size-spectra have been sampled at various locations and represented in many ways. Individual size may be in units of length, volume, wet or dry mass, and counted over an area or volume, which may be transformed into biomass or energy density. Individuals may be binned into any number of logarithmic or linear size-classes, or represented as a probability or cumulative density function, and fit using various methods such as least squares or maximum likelihood (*2, 4, 6*). These varied published representations can obscure commonalities among observed patterns, which, when expressed as a size-frequency distribution, tend to be strikingly similar with exponents near a value of −1, as revealed by a recent meta-analysis (*4*), and shown in Table S3 and Fig. S2.

**Fig. S2.**
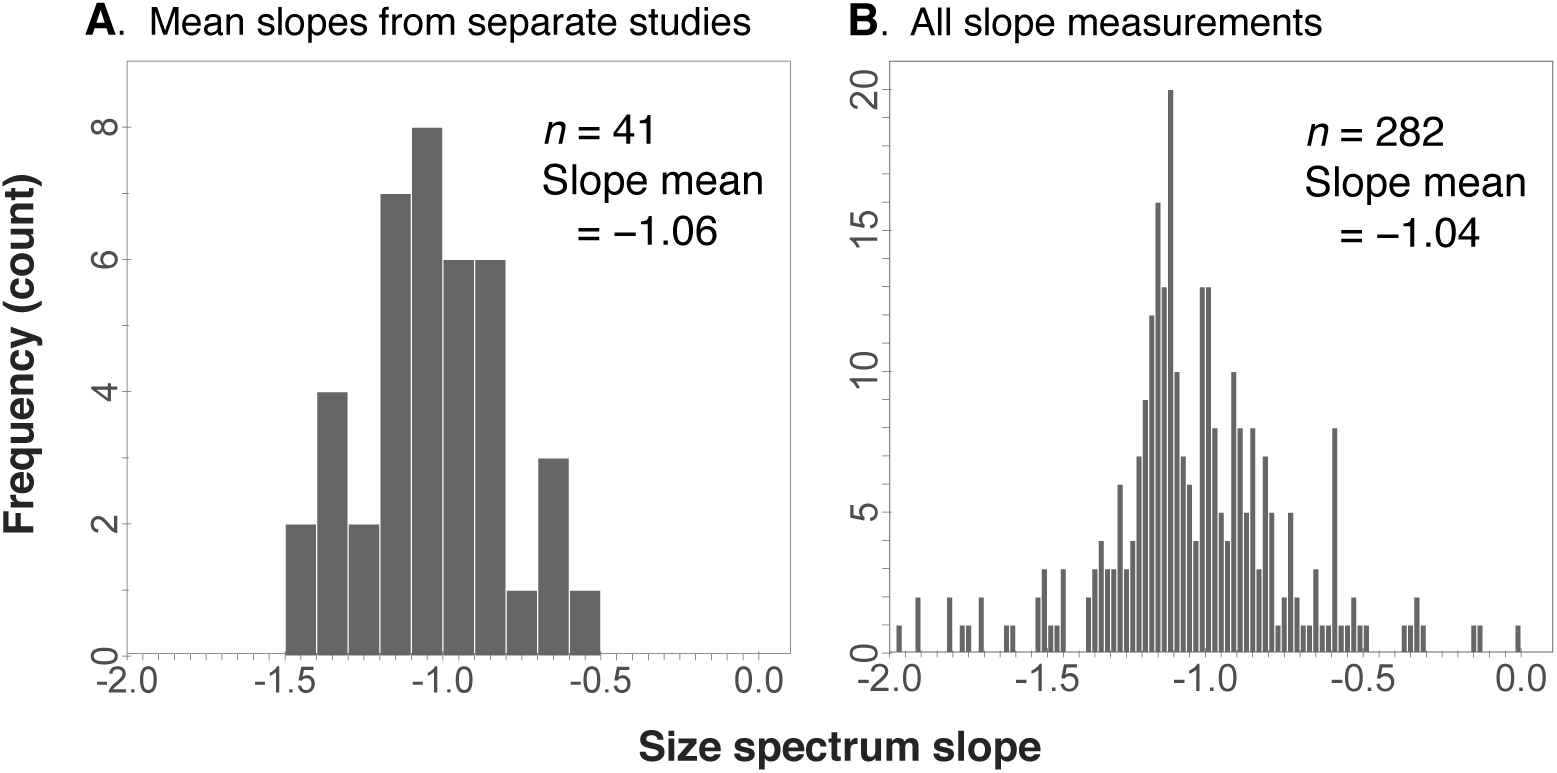
Summary of size spectra slopes from the literature. These slopes represent the normalized biomass spectra slopes, or equivalently, the number of individuals in each logarithmic size class. **A.** A histogram of the mean slope from each of 41 separate studies. **B.** A histogram of all 282 slopes reported across 41 studies. All data are from (*4*), and reproduced in the Supplementary Data file, with some of the more important studies listed in Table S3.

**Table S3.**
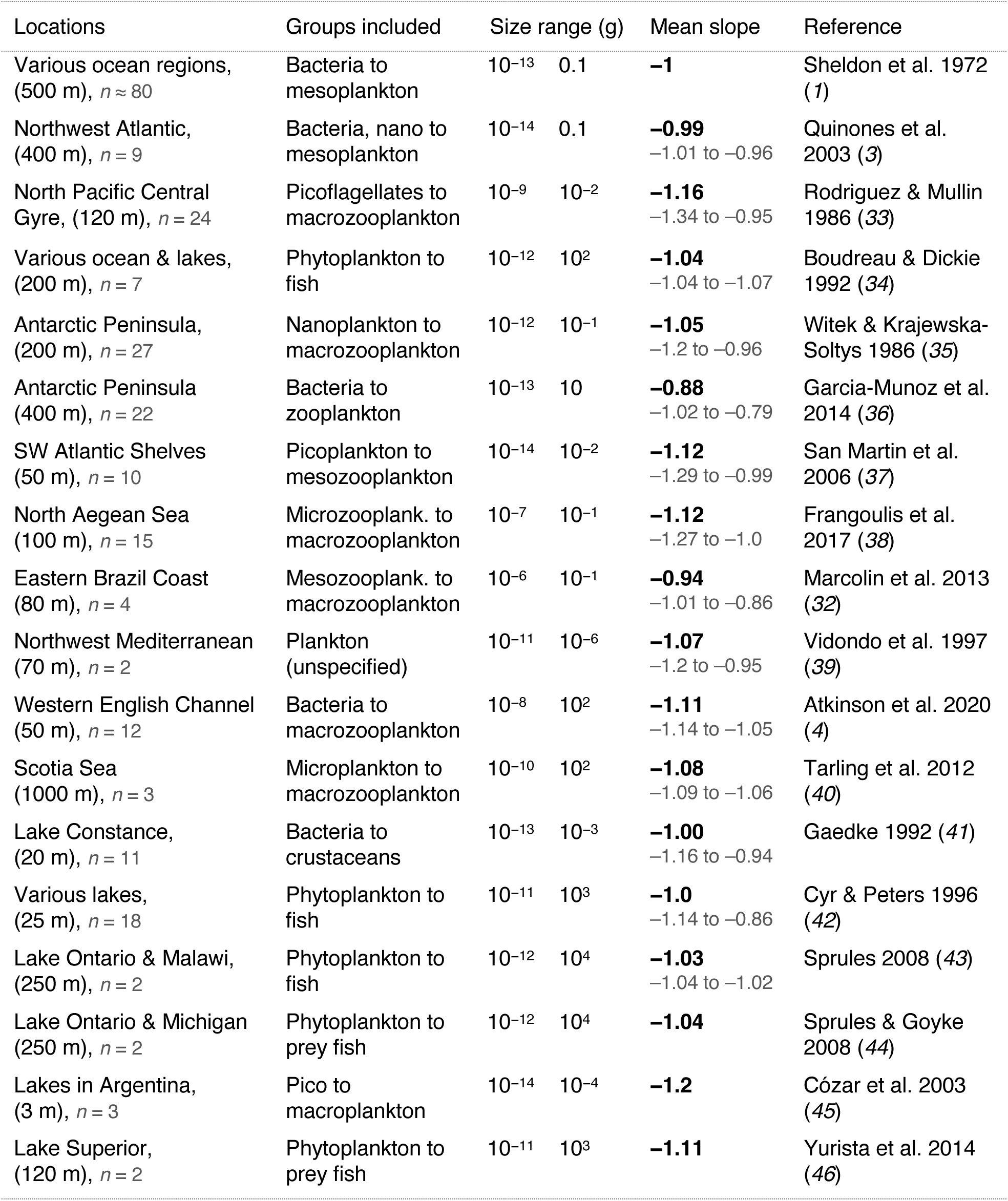
Summary of some important size spectra studies. The number of locations (*n*) may refer to separate regions or distinct time periods (with maximum depth shown in meters; Sheldon 1972 included a few size spectra greater than 500 m, but most were <200m). The reported smallest to largest groups and order of magnitude range of body sizes (grams fresh mass) is approximated. The mean slope of the size-spectrum (normalized biomass) is reported with the range of slope values underneath, where available. Further studies are listed in (*4, 32*).

| Locations | Groups included | Size range (g) |  | Mean slope | Reference |
| --- | --- | --- | --- | --- | --- |
| Various ocean regions, (500 m), *n* ≈ 80 | Bacteria to mesoplankton | 10<sup>-13</sup> | 0.1 | **-1** | Sheldon et al. 1972 (1) |
| Northwest Atlantic, (400 m), *n* = 9 | Bacteria, nano to mesoplankton | 10<sup>-14</sup> | 0.1 | **-0.99**<br>-1.01 to -0.96 | Quinones et al. 2003 (3) |
| North Pacific Central Gyre, (120 m), *n* = 24 | Picoflagellates to macrozooplankton | 10<sup>-9</sup> | 10<sup>-2</sup> | **-1.16**<br>-1.34 to -0.95 | Rodriguez & Mullin 1986 (33) |
| Various ocean & lakes, (200 m), *n* = 7 | Phytoplankton to fish | 10<sup>-12</sup> | 10<sup>2</sup> | **-1.04**<br>-1.04 to -1.07 | Boudreau & Dickie 1992 (34) |
| Antarctic Peninsula, (200 m), *n* = 27 | Nanoplankton to macrozooplankton | 10<sup>-12</sup> | 10<sup>-1</sup> | **-1.05**<br>-1.2 to -0.96 | Witek & Krajewska-Soltys 1986 (35) |
| Antarctic Peninsula (400 m), *n* = 22 | Bacteria to zooplankton | 10<sup>-13</sup> | 10 | **-0.88**<br>-1.02 to -0.79 | Garcia-Munoz et al. 2014 (36) |
| SW Atlantic Shelves (50 m), *n* = 10 | Picoplankton to mesozooplankton | 10<sup>-14</sup> | 10<sup>-2</sup> | **-1.12**<br>-1.29 to -0.99 | San Martin et al. 2006 (37) |
| North Aegean Sea (100 m), *n* = 15 | Microzooplank. to macrozooplankton | 10<sup>-7</sup> | 10<sup>-1</sup> | **-1.12**<br>-1.27 to -1.0 | Frangoulis et al. 2017 (38) |
| Eastern Brazil Coast (80 m), *n* = 4 | Mesozooplank. to macrozooplankton | 10<sup>-6</sup> | 10<sup>-1</sup> | **-0.94**<br>-1.01 to -0.86 | Marcolin et al. 2013 (32) |
| Northwest Mediterranean (70 m), *n* = 2 | Plankton (unspecified) | 10<sup>-11</sup> | 10<sup>-6</sup> | **-1.07**<br>-1.2 to -0.95 | Vidondo et al. 1997 (39) |
| Western English Channel (50 m), *n* = 12 | Bacteria to macrozooplankton | 10<sup>-8</sup> | 10<sup>2</sup> | **-1.11**<br>-1.14 to -1.05 | Atkinson et al. 2020 (4) |
| Scotia Sea (1000 m), *n* = 3 | Microplankton to macrozooplankton | 10<sup>-10</sup> | 10<sup>2</sup> | **-1.08**<br>-1.09 to -1.06 | Tarling et al. 2012 (40) |
| Lake Constance, (20 m), *n* = 11 | Bacteria to crustaceans | 10<sup>-13</sup> | 10<sup>-3</sup> | **-1.00**<br>-1.16 to -0.94 | Gaedke 1992 (41) |
| Various lakes, (25 m), *n* = 18 | Phytoplankton to fish | 10<sup>-11</sup> | 10<sup>3</sup> | **-1.0**<br>-1.14 to -0.86 | Cyr & Peters 1996 (42) |
| Lake Ontario & Malawi, (250 m), *n* = 2 | Phytoplankton to fish | 10<sup>-12</sup> | 10<sup>4</sup> | **-1.03**<br>-1.04 to -1.02 | Sprules 2008 (43) |
| Lake Ontario & Michigan (250 m), *n* = 2 | Phytoplankton to prey fish | 10<sup>-12</sup> | 10<sup>4</sup> | **-1.04** | Sprules & Goyke 2008 (44) |
| Lakes in Argentina, (3 m), *n* = 3 | Pico to macroplankton | 10<sup>-14</sup> | 10<sup>-4</sup> | **-1.2** | Cózar et al. 2003 (45) |
| Lake Superior, (120 m), *n* = 2 | Phytoplankton to prey fish | 10<sup>-11</sup> | 10<sup>3</sup> | **-1.11** | Yurista et al. 2014 (46) |

Another consideration with respect to the Sheldon hypothesis is that the “pristine” biomass distribution, which existed prior to human hunting and habitat modification, differs from the current biomass distribution. On the one hand, the interest in the Sheldon hypothesis as a possible universal law-like property of intact aquatic systems would prompt use of the reconstructed pristine biomass, which we have done here. On the other hand, it is of significant interest to know how human impacts might have altered the size-spectrum through a long history of harvesting, as well as the current expectations of how humans may impact the size-spectrum through climate change (*12*). We therefore also provide an estimate of how historical impacts from fishing and whaling have shaped the current size-spectrum, as well as projecting changes in biomass under a pessimistic scenario (RCP8.5) of future changes in climate from year 2000 to 2100 (Fig. 3).

A number of assumptions must be made in order to aggregate individuals into body mass size-classes. We chose the major groups of bacteria, phytoplankton, zooplankton, fish and mammals, as these capture common distinctions between functional groups, but more importantly, they demarcate important differences in our methods for estimating biomass, as described further in the section “Data and methods”. Among some major groups, further sub-divisions were necessary for improving our estimates given the data available (listed in Table S2). Finally, these sub-divided groups were split into order-of-magnitude size classes. While admittedly coarse, this represents a common level of resolution that can be applied to all major groups with similar confidence. For many groups, the available data allows much more refined size-class binning, but we instead set the bin size to a coarse order of magnitude to avoid making additional assumptions for less resolved group size distribution. This provided us with 23 separate and relatively well estimated data points on which to construct a regression, which is discussed further in “Statistical considerations”.

#### Uncertainty

Uncertainties in our estimates come from a range of sources. We estimate uncertainty for each major group as a multiplicative fold factor, which in turn determines our global estimate of the shape of the size-spectrum frequency distribution.

##### Measurement error

Estimation of phytoplankton biomass depends on satellite estimates of chlorophyll a concentration at the surface, while estimates of bacteria and zooplankton are based on interpolations from hundreds of thousands of sample estimates. These estimates depend on well calibrated observing instruments. Small drifts in instrument calibration can cause significant errors in reported chlorophyll concentration, and abundance measures, which in turn drive errors in estimates of group biomass (*47*). Zooplankton biomass samples have been taken with dozens of different gear types and mesh sizes over the past 60 years (*48*). However, many of these gear and mesh types are not suitable to capture small or large zooplankton due to passive or active avoidance, aggregation and diurnal migration, leading to (usually) underestimates of total zooplankton biomass (*49*). Further, the ability of many mesopelagic fish to avoid nets can produce taxon-specific biases that are difficult to estimate (*50*). Because of the inherent difficulties with scientific sampling, some fish estimates make use of fishery data to help constrain biomasses. Industrial fishing activities capture a large fraction of global fish production, providing independent constraints on coupled fish-fishery models (*13*), though they also have large uncertainties due to unreported, underreported and illegal catches (*51*). Finally, in the case of mammals, most abundance estimates come from visual observation at the sea surface or in breeding grounds. In addition to the measurement errors associated with different methods, there is also uncertainty in how comparable these different methodologies are (*17*). This is particularly relevant to our estimates of uncertainty for each group and size class.

##### Sampling bias

The ocean covers 2/3 of the planet, and large parts of it have not been sampled thoroughly. What is more, the environment is continually fluctuating, which can cause large temporal variability at any point in the ocean. As a result, there is the potential for large biases in the spatial and temporal distribution of observational samples (*52*) which is likely to greatly exceed the measurement error. We address this bias using statistical and/or process-based models that capture how biomass abundance varies with the environment, and then resample the models at the global scale using realistic average environmental conditions. Although the models have large uncertainties for predicting the biomass under any given set of environmental conditions, they have much less uncertainty associated with the global mean values, which are the target of interest here.

##### Major group size distributions

Much research over the past 50 years has revealed that, within limited taxonomic domains, the size-frequency distribution is characteristically the same as the broader size-spectrum that we show in Fig. 1 (power law distribution with an exponent near −1; see Table S3). For some major groups, however, very little is known of the size distribution, which is particularly true for bacteria, for which we are aware of only a single study that explicitly studied their size distribution in the open ocean (*53*). We therefore test this uncertainty by using alternative size distributions and undertaking a sensitivity analysis of our assumption of an equal distribution of biomass across order of magnitude size bins for each major group, described below in the section “Group size distributions and sensitivity analysis”.

##### Boundaries of the Sheldon hypothesis

There is uncertainty regarding the physical and taxonomic boundaries of what ought to delimit the “Sheldon hypothesis” in the ocean. Sheldon first hypothesized that a constant biomass across log size classes extended from bacteria to whales, with little indication of the physical boundaries or the many taxonomic groups, in which this pattern should be relevant (*1*). For example, should we assume that the physical boundary in which the pattern is most relevant is only the open surface waters of the epipelagic, or might it extend to the mesopelagic or all the way to the seafloor and seafloor sediment. Should the physical environment include benthic and demersal species, brackish waters and mangrove ecosystems, etc.? Likewise, are we to assume that all taxonomic groups whose niche overlaps with these environments should be included in the Sheldon hypothesis, or are we to include only species that physically reside in these environments, excluding say, seabirds and polar bears? While the inclusion/exclusion of minor taxonomic groups should have little bearing on the overall Sheldon spectrum, the physical boundary, and particularly the depth to which we delimit the problem alters the shape of the distribution. It remains questionable whether a formal test of the Sheldon hypothesis from bacteria to whales is appropriate down to the sea floor, where few major groups exist, thus extending the boundary of the problem to the group with the largest extent, far beyond the existence of most other groups. It also becomes problematic with comparisons to previous size spectra studies, which do not extend to such depths (Table S3 lists size spectra studies which typically do not extend beyond 500 meters). Nonetheless, we address this important uncertainty by showing the results for two domains in the ocean: the upper 200m and the full water column (see Fig. 2 and Table S2).

##### Calculating uncertainty bounds

For each major group, uncertainties are represented as 95% confidence intervals of the average biomass estimate (Fig. 2 A). Following the approach of (*15*), we report these uncertainty ranges as multiplicative fold changes (×/÷) from the mean, rather than additive changes (±). This is done because the distribution of sample data is best approximated by a log-normal, and the geometric mean (i.e. the average on log scale) will give an estimator that is more robust to outliers, particularly when data are sparse, or are biased towards particular geographic regions or taxonomic groups. Multiplicative uncertainty is more robust to large fluctuations and appropriate for log-normally distributed data. Moreover, much of the error in estimation, as well as the natural fluctuations in time or in space or among taxa, are multiplicative rather than additive. Given the range of sources we use to calculate our biomass estimates, the method we used to estimate uncertainty was not the same for all groups. The extent to which uncertainty bounds across methodologies can be compared remains difficult to evaluate. We summarize the method used to estimate uncertainty for each major group below.

In addition to these general uncertainties, there are also uncertainties associated with each major group. Phytoplankton are estimated globally using satellite images of surface chlorophyll, which can be biased in converting from ocean color to total depth-integrated biomass. Heterotrophic bacteria and zooplankton are estimated from water samples, but their biomass varies spatially and data coverage is patchy (mapped points in Fig. 1 A). Fish aggregate, migrate and can escape capture, so we rely on more comprehensive commercial harvest data to constrain two independent global process models, which can be biased by assumptions. And large marine mammals can individually range over the entire ocean, requiring global population estimates for many species that are necessarily incomplete. Finally, we seek pristine estimates of fish and mammals, prior to commercial exploitation, requiring careful consideration of human impacts over the past two centuries. These uncertainties are discussed further below and in greater detail in the section “Data and methods”.

**Table S4.**
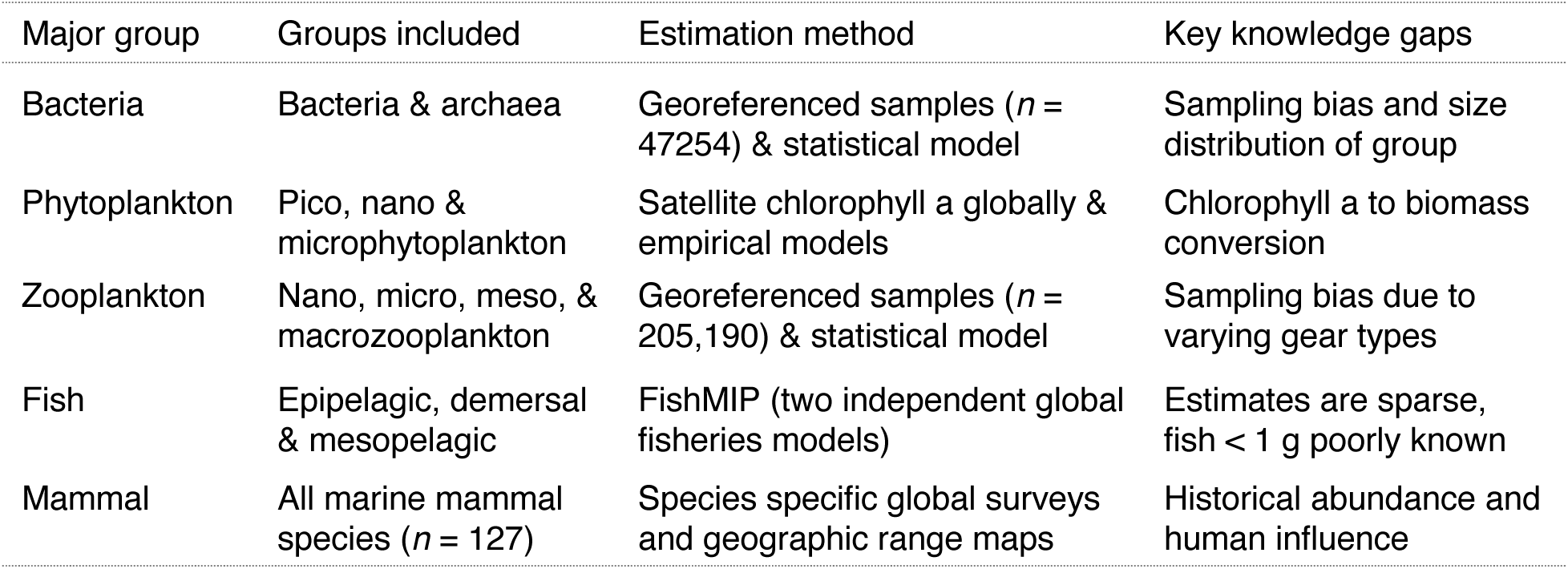
Summary of estimation methods by major groups and key gaps in knowledge.

##### Bacteria and zooplankton

Global estimates of abundance for bacteria and biomass for all zooplankton groups (nano-, micro-, meso-, and macro-zooplankton, as well as all other animals that pass through a zooplankton life-stage) were calculated using generalized linear models fit to log-transformed sample measurements. For our statistical models of bacteria and zooplankton we follow the approach of (*15, 16*) in reporting our uncertainties using the geometric mean of two measures of uncertainty: 1) the standard error from 1000 bootstrap predictions of mean global abundance or biomass from the statistical model, and 2) the standard deviation of the log-transformed sample data on which the statistical model was fit. The geometric mean of the two uncertainty measures is then multiplied by 1.96 to obtain the 95% confidence interval for the mean estimate in log-space. We use the geometric mean of these two uncertainty measures to balance their limitations. For the first uncertainty estimate, the standard error of the global mean biomass estimate will be smaller the better the predictive capability of our statistical model. However, this uncertainty does not consider the presence of bias or measurement error in the sample data used to fit the statistical model, therefore this uncertainty estimate is most likely an underestimate. On the other hand, the standard deviation of the log-transformed data would be an overestimate of the uncertainty of average biomass, given it does not take into account the decrease in uncertainty that comes from the averaging that takes place when the statistical model is fit.

For bacteria, an additional step is required where mean global abundance is multiplied by average bacteria cell size to estimate mean global biomass. We incorporate this added uncertainty in our global bacteria biomass estimate by multiplying 95% confidence interval of geometric mean bacteria cell size with the 95% confidence interval of the geometric mean global abundance of bacteria.

##### Phytoplankton

Global phytoplankton biomass was calculated from satellite surface chlorophyll observations. We used an empirical equation from (*54*) to derive total chlorophyll in the water column from satellite surface estimates, and then another empirical equation from (*55*) to convert total chlorophyll into phytoplankton carbon and finally wet biomass. These empirical equations were published with reported 95% confidence intervals in log-space, and we obtained the cumulative 95% confidence interval by multiplicatively combining the reported 95% confidence intervals from these two empirical equations.

##### Fish

We split ‘fish’ (which includes large cephalopds and invertebrates) into two groups; mesopelagics and non-mesopelagics. For mesopelagic fish, we quantified the uncertainty of our estimate using results from (*56*), who assessed how uncertainty in the composition of the mesopelagic community causes uncertainty in estimates derived from bioacoustics surveys. For non-mesopelagic fish, including cephalopods and benthic invertebrates, we obtained estimates of global biomass from two process-based marine ecosystem models; the FishErIes Size and functional Type model (FEISTY; (*14*)) and the BiOeconomic mArine Trophic Size-spectrum model (BOATS; (*13*)). We used publicly available output from these models to derive our global biomass estimates (both models’ output available at https://www.isimip.org). To calculate the uncertainty range for our estimates of non-mesopelagic fish biomass, we follow the approach of (*16*), who took the 90% confidence interval of global fish biomass from (*29*) as representative of the uncertainty of their non mesopelagic fish biomass estimate.

##### Mammals

Estimates of marine mammal uncertainty is obtained from additional species population data from multiple additional sources. These include current mammal abundance estimates listed on IUCN species pages, as well as data drawn from Appendix 4 of (*28*), which lists minimum, mean and maximum values for 115 of 126 marine mammal species. The vast majority of these estimates tend to be within a multiplicative factor of three of our current values. Obtaining uncertainty estimates for pre-whaling abundance is more challenging, but are seldomly more than a factor of ten different from other estimates, and on average are a factor of 4.3 of maximum current values from (*28*). Pooling all estimates including those of IUCN (*57*) and current values of (*28*) (which may not be independent of one another) gives a mean fold uncertainty of 2.2 (following methods described in (*15, 16*)). Given that our knowledge of the pre-whaling period is necessarily vague, we considered a fold uncertainty value of 5 to be cautious, and include all additional abundance estimates used for estimating uncertainty in the Supplementary Data file.

Further information about uncertainty are described under major group headings in the section “Data and methods” below.

### Data and methods

In this section, we describe our data sources, methods and models for estimating global ocean abundance and biomass, as well as procedures for estimating uncertainty in these values.

#### Bacteria

There were two steps to obtain our estimate of global bacteria biomass. First, we fit a statistical model of bacteria abundance with monthly georeferenced chlorophyll a, sea-surface temperature and 1° bathymetry data to interpolate global bacteria abundance. These environmental variables were taken from monthly climatologies of MODIS-Aqua (Moderate Resolution Imaging Spectroradiometer aboard the Aqua spacecraft) 4km measurements, from 2002-2016. Similarly, 1° bathymetry data was used to calculate the depth-integrated biomass of bacteria in each 1° region of the global ocean. We then used *in situ* samples of individual bacteria cell size from coastal and open ocean sites to estimate the average individual bacteria cell size, as well as the size range of bacteria. We then multiplied our global bacteria abundance estimate by our estimate of average bacteria cell size to obtain global bacteria biomass. Our data sources for bacteria abundance did not distinguish between bacteria and archaea, so we include both of these groups in our bacteria biomass estimate.

##### Data sources of bacteria abundance

1. 5671 observations from Wigington et al. 2016 (*58*) https://doi.org/10.1038/nmicrobiol.2015.24
2. 39767 observations from Buitenhuis et al. 2012 (*59*) https://doi.pangaea.de/10.1594/PANGAEA.779142
3. 1816 observations from Lara et al. 2017 (*60*) http://doi.org/10.1126/sciadv.1602565

##### Statistical model of bacteria abundance

We fit a generalised linear model, with log_10_ bacteria abundance as the response variable, with a linear relationship for the environmental predictors log_10_ chlorophyll, temperature, and a third order polynomial for log_10_ sample depth (Figure 1). All predictors are significant, and the model has an R^2^ of 74%.

##### Total global bacteria abundance

For each grid square, we use temperature, bathymetry and chlorophyll to calculate the depth-integrated density of bacteria (# m^2^). We then summed all grid squares to obtain a total global abundance of **1.41 x 10*^29^*** bacteria (×/÷3.4-fold 95% confidence interval). This number is close to Bar-On et al. 2019 (*16*) estimate of 1.2 x 10^29^, which was derived from three previous studies of total bacteria abundance (*61*): 8 x 10^28^; (*62*): 1.7 x 10^29^; (*59*): 1.2 x 10^29^).

**Fig. S3.**
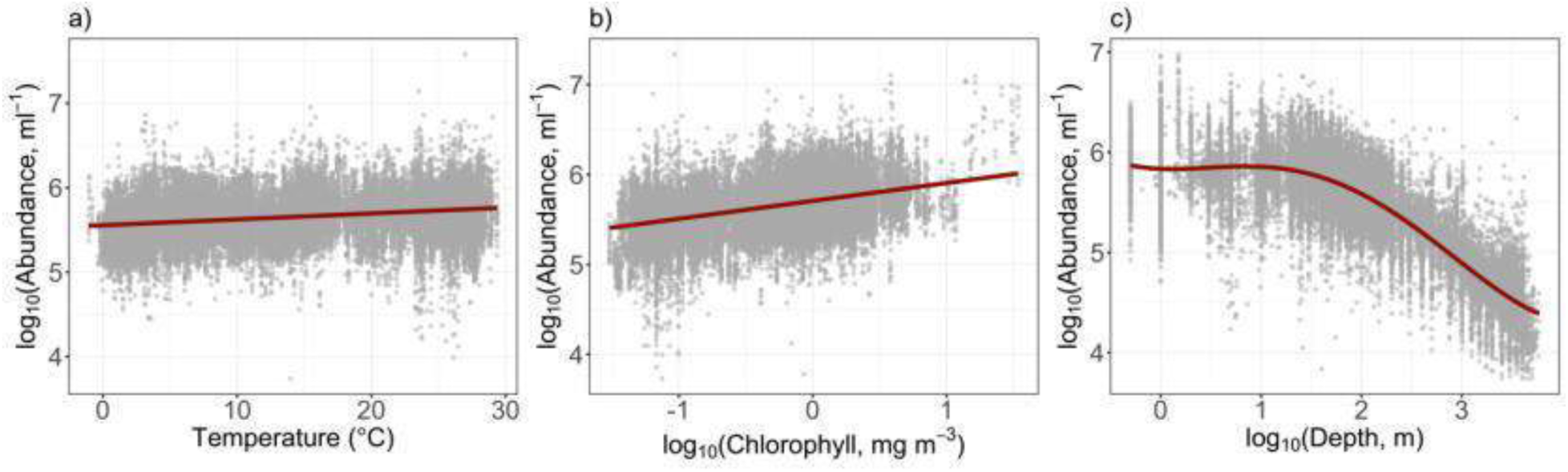
Model fit to each of the predictor variables for bacteria abundance, with partial residuals.

##### Mean individual bacteria wet weight

We obtained individual bacteria cell size observations (both carbon and wet weight) from five studies where *in situ* samples were taken from pelagic and coastal regions (*63–67*). For the two largest studies ((*63*): *n* = 145, Sargasso Sea) and ((*64*): *n* = 201, coastal Norway and Denmark), individual bacteria were measured with an electron microscope, after microplankton cells containing chlorophyll were identified and removed. Carbon content was determined using X-ray microanalysis, calibrated on particles with known chemical composition. The cell size distributions (in wet weight) for these studies are similar to distributions from other studies (*53*,*68*) that used flow cytometry—considered a reference technique in bacteria community analysis (*69*)—to count and measure individual bacteria. Contrary to (*70*), we didn’t find a statistically significant difference between the distribution of open ocean – Sargasso Sea – observations (95% CI for mean = 7.2-11.6 fg C), and combined coastal observations (95% CI for mean = 9.1-11.4 fg C), so we combined both coastal and open ocean data together, and assumed that individual bacteria cell sizes across the global ocean can be well approximated by a single lognormal distribution.

For each sample used to build the distribution of individual bacteria size in carbon, we also obtained wet weight (biovolume). Bacteria wet weight observations also follow a lognormal distribution (Fig. S4) with mean individual bacteria wet weight of 0.109 pg (95% confidence interval – 0.096, 0.124 pg; ×/÷ 0.14-fold uncertainty).

**Fig. S4.**
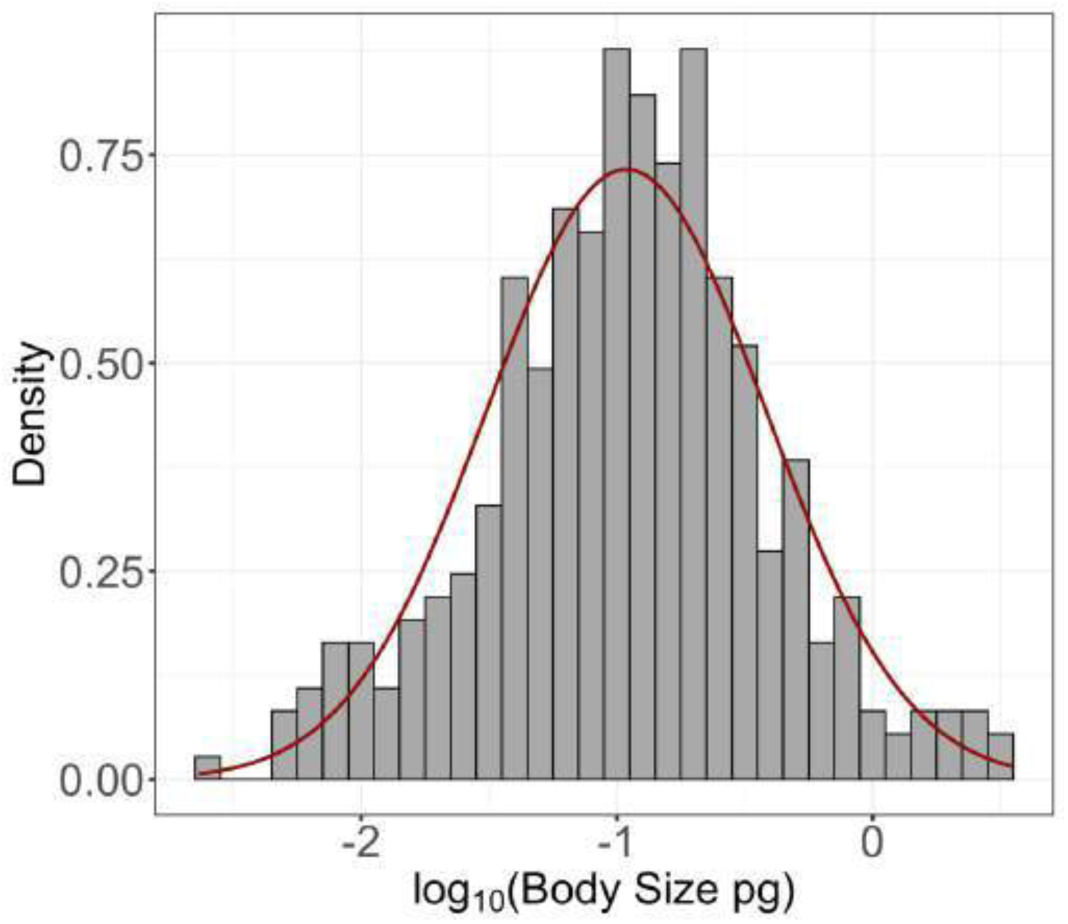
Histogram of bacterial cell sizes in log_10_ pg wet weight.

##### Global bacteria biomass and size range

Using our mean wet weight bacteria cell size of 0.109 pg, we calculate global bacteria biomass is **14.48 Pg** (cumulative 95% confidence interval is ×⁄÷ 3.85-fold). We used the 99% confidence interval of our bacteria cell size distribution (Figure 2) to define the size range of bacteria for this study to be 10^-2.3^ – 10^0.5^ pg.

#### Phytoplankton

To estimate global phytoplankton biomass, we used satellite chlorophyll observations from MODIS-Aqua, averaged over 2002-2016, and empirical equations of chlorophyll with depth and by functional type, to calculate the global biomass of pico, nano and microphytoplankton. Our estimate of total phytoplankton biomass includes both autotrophs and mixotrophs, since both of these groups contain chlorophyll and so are represented in satellite estimates.

##### Chlorophyll a from pico, nano and microphytoplankton

First, we calculate the percentage of surface chlorophyll from pico (< 2%m), nano (2-20%m) and microphytoplankton (20-200%m) with the empirical model developed by (*71*). The model was developed with a dataset of phytoplankton pigments from the Atlantic Ocean, and shows close agreement with other global models of phytoplankton functional types developed with datasets for different regions (Figure 4. 3 of IOCCG Report, (*72*)). We assume the percentage of chlorophyll from each group is the same as the surface throughout the entire water column. This assumption is supported by Figure 4. 3 in the IOCCG (*72*), which shows that the contribution of different phytoplankton groups from the chlorophyll a across the entire euphotic zone (*54*) is in close agreement with the contribution from the Brewin model (*71*), which used only surface chlorophyll a.

##### Chlorophyll a in euphotic zone

To calculate total chlorophyll a in the euphotic zone from surface chlorophyll a, we use the empirical equations in (*54*). Uitz et al., (*54*) gives two equations for total chlorophyll a from surface chlorophyll a, one for stratified waters (defined as water where euphotic zone depth is less than the mixed layer depth) and another for mixed waters (defined as water where euphotic zone depth is greater than mixed layer depth). To apply Uitz’s equations, we divide the global ocean into mixed and stratified grid squares, using average euphotic zone depth for 2002-2019 from Modis-AQUA, derived with the Lee algorithm (*73*), and annual average mixed layer depth derived from Argo profiles (*74*).

The Uitz equations (*54*) are fit to a sample set from 2419 stations across the global ocean, with ∼1800 from stratified waters and ∼600 from mixed waters. The fit of the Uitz equations to the data is close to that of earlier work from (*75*), and was chosen for its use of an extensive dataset to derive total chlorophyll a in the euphotic zone from satellite output. In terms of uncertainty, there was a 1.4-fold 95% prediction interval in the relationship between surface chlorophyll a and total chlorophyll a in the euphotic zone (*54*).

##### Conversion to wet biomass by size class

To convert total chlorophyll a concentration to carbon, we used the empirical equation from (*55*), who fit a relationship between chlorophyll and phytoplankton biomass (carbon concentration), with surface observations from coastal and open ocean stations across the global ocean. The linear relationship between log chlorophyll and log carbon concentration in (*55*), had an R^2^ value of 0.9 and a 2.3-fold 95% prediction interval. In each grid square, to calculate total carbon content in the water column, we divided total chlorophyll from step 2 by euphotic zone depth and then used this average chlorophyll concentration to calculate the average carbon concentration with the Marañón equation (*55*). We then calculated average total carbon content in the water column in each grid square by multiplying average carbon concentration by the euphotic zone depth. We then used the percentages calculated in step 1 to break total carbon into pico, nano and microphytoplankton groups. Finally, to convert total carbon of each group into wet weight, we assumed a wet weight to carbon ratio of 10:1 (*34*).

##### Individual cell size ranges converted to wet weight

The size ranges for pico, nano and microphytoplankton are typically expressed in um ESD: 0.2μm – 2 μm for picophytoplankton, 2 μm – 20 μm for nanophytoplankton and 20 μm – 200 μm for microphytoplankton (*71*). We convert these cell size ranges to wet weight, assuming 1 g wet weight is equal to 1cm^3^.

We calculated global picophytoplankton biomass to be 2.15 Pg, nanophytoplankton biomass 1.8 Pg and microphytoplankton biomass to be 1.35 Pg. Altogether, total phytoplankton biomass **is 5.3 Pg** (cumulative 95% confidence interval is ×⁄÷ 3.2).

#### Zooplankton

To interpolate sample estimates of zooplankton biomass, we used satellite chlorophyll a, bathymetry and sea-surface temperature as environmental predictor variables. Monthly chlorophyll a and sea-surface temperature measurements were appended to observations, based on the month in which they were taken, as well as their latitude and longitude. These environmental variables were taken from monthly climatologies of MODIS-Aqua 4km measurements, from 2002-2016. Similarly, 1° bathymetry data was used to calculate the depth-integrated biomass of zooplankton in each 1° region of the global ocean.

Zooplankton can be broken into four size ranges: nano (0.8µm −5µm), micro (5µm - 200µm), meso (200µm-2cm), macro (0.2-10cm) (*17, 76*). Nano and microzooplankton cover protists groups such as heterotrophic flagellates, dinoflagellates, ciliates and juvenile mesozooplankton (*77, 78*). Mesozooplankton cover groups such as copepods, larvaceans, amphipods and giant rhizaria (*79, 80*). Macrozooplankton include groups such as chaetognaths, euphausiids, tunicates, fish larvae, ctenophores and cnidaria (*81*).

##### Nanozooplankton

For this group, we were unable to find reliable resources with which to generate a global statistical model, so we used the estimate of (*16*) of **2.5 Pg** wet weight biomass (assuming a 10-1 wet weight to carbon conversion) for this group. No uncertainty bound was provided for this estimate, as it is derived from only one source (*78*), so we use the ×∕÷ 6.5-fold 95% confidence interval calculated for microzooplankton (see below), which overlaps taxonomically with the nanozooplankton (*77, 78*).

##### Microzooplankton

Data derived from 4579 observations from Buitenhuis et al. 2010 (*77*). https://doi.org/:10.1029/2009GB003601

After applying quality control to the data (e.g., filtering NA observations for biomass, depth sampled) we were left with 3866 observations. For the 46 observations where depths are greater than 1000m, depth was set to 1000m.

We used a generalised linear model, with a linear relationship between log_10_ zooplankton biomass and temperature, a third order polynomial for log_10_ sample depth and another third order polynomial for log_10_ chlorophyll. All predictors are significant, and the model has an R^2^ of 27%. Although the R^2^ is low, the relationship of log biomass with environmental variables is reasonable (Fig. S5). For instance, the chlorophyll relationship is not uniformly increasing, but is highest in mesotrophic waters, which matches the results of the model in (*77*).

**Fig. S5.**
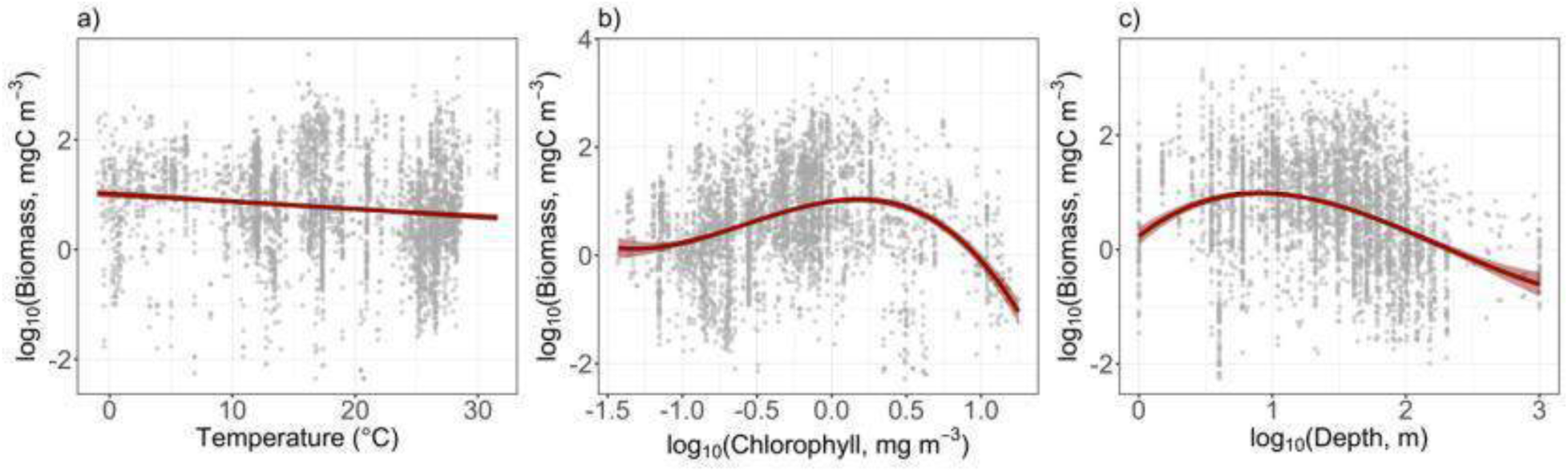
Model fit to each of the predictor variables for microzooplankton, with partial residuals.

For each grid square, we use temperature, bathymetry and chlorophyll, and a carbon to wet weight conversion factor of 10 (*34*) to calculate the depth-integrated wet weight biomass of microzooplankton (g m^3^) over the water column in each grid square, to obtain a total global biomass of **4.79 Pg** (95% confidence interval for this estimate is ×/÷ 6.5-fold).

##### Mesozooplankton

Mesozooplankton are not only comprised of groups such as meroplanktonic larvae and copepods, but also giant protists from the group Rhizaria. A recent study (*79*) indicated that although giant Rhizaria are a major fraction of the mesozooplankton, their delicate nature means that they are heavily undersampled in conventional sampling techniques such as the ones used to build the COPEPOD database. To address this, we obtained an estimate of global giant rhizaria biomass separately (see next sub-section) to ensure that this group was properly incorporated into our overall estimate of mesozooplankton biomass. Here we describe all other mesozooplankton biomass estimates.

Data derive from 214,922 observations from the COPEPOD database (*48*): https://www.st.nmfs.noaa.gov/copepod/atlas/html/taxatlas_b400.html.

After applying quality control to the data (e.g., filtering NA observations for biomass, volume filtered, depth sampled, removing samples from mesh sizes less than 200µm and greater than 2000µm) we were left with 176,000 observations.

We used a generalized linear model, with a linear relationship between log_10_ zooplankton biomass, and log_10_ chlorophyll, temperature, and a third order polynomial for log_10_ sample depth. Observations across measurement types were not converted to carbon, so we also included a factor variable for the measurement type (Fig. S6). All predictors are significant, and we get an R^2^ of 83% for this model. The majority of this variance explained comes from the measurement type factor, and when that factor is excluded the R^2^ is 18%. The relationships with the environmental variables and sample depth is reasonable, as is the measurement type factor – with wet weight biomass having a higher mean value than the other methods. The spread of the residuals is very wide, about two orders of magnitude around the predicted values (Fig. S6).

**Fig. S6.**
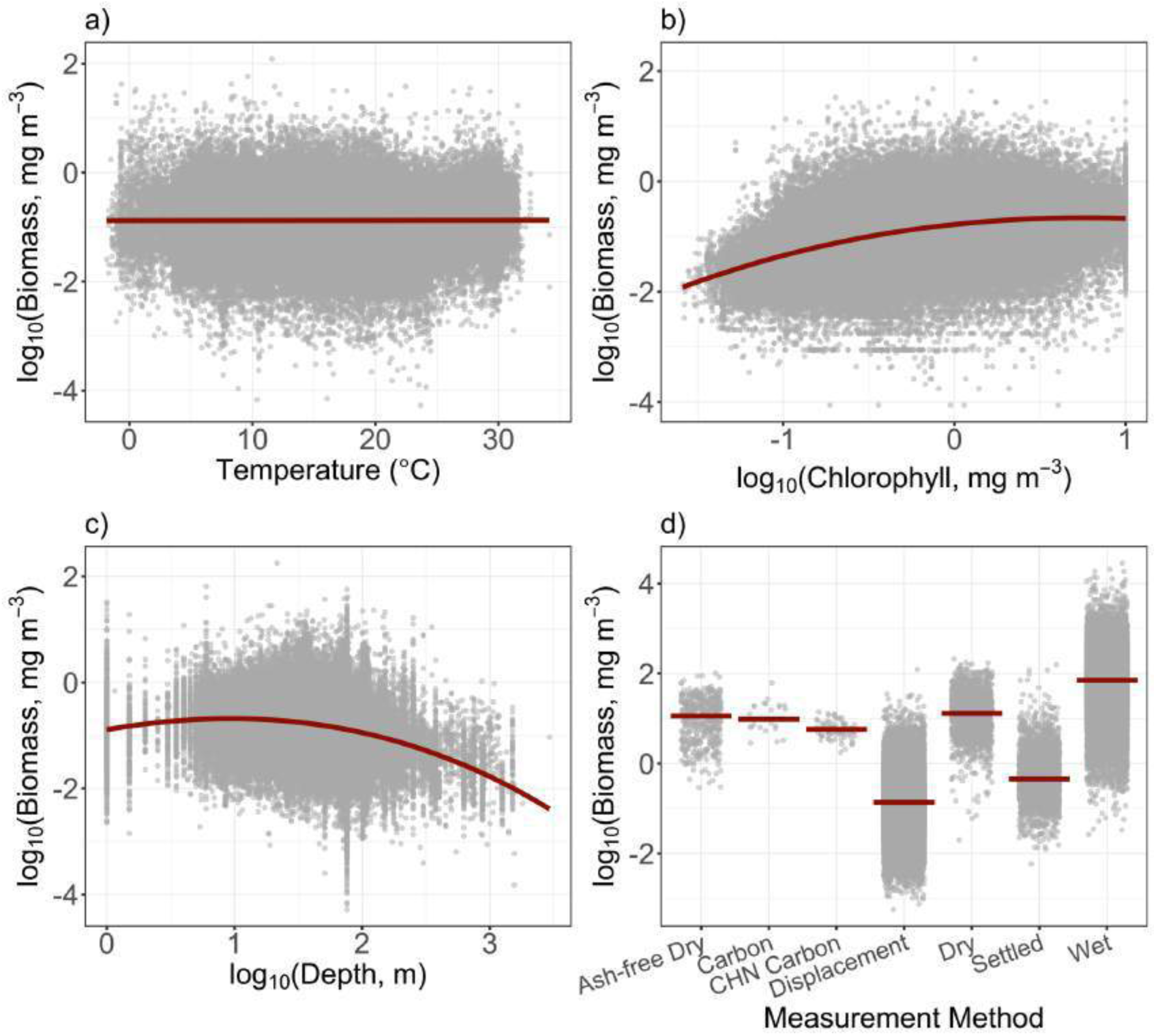
Model fit to each of the predictor variables for mesozooplankton, with partial residuals.

##### Giant rhizaria

Data derive from 1509 observations from the supplement to Biard et al. 2016 (*79*). https://doi.pangaea.de/10.1594/PANGAEA.858156

Each observation gives the total integrated carbon biomass mg C m^-2^ over a transect of the water column, with 681 observations covering 0-200m, and the remaining 828 observations covering 0-500m. We fit a statistical model to the 0-200m and 0-500m datasets separately, using the 0-500m dataset to estimate total giant rhizaria biomass over the water column (following the approach of (*16*)). Although cell abundance for protists is much lower across the deep ocean (*82*), the lack of sampling below 500m means that total rhizaria biomass may be underestimated.

For our estimate of giant rhizaria biomass over the total water column, and from 0-200m, we used separate generalized linear models, each with a second order polynomial relationship between log_10_ rhizaria biomass (from 0-500m or 0-200m), and log_10_ chlorophyll and temperature (Fig. S7). Both temperature and log_10_ chlorophyll predictors are significant for both models, and we get an R^2^ of 25.9% for the model fitted to 0-200 m rhizaria biomass samples, and an R^2^ of 22.7% for the model fitted to 0-500 m samples. The fitted relationships between log_10_ chlorophyll, temperature and log_10_ biomass were similar across both models (Fig. S7).

**Fig. S7.**
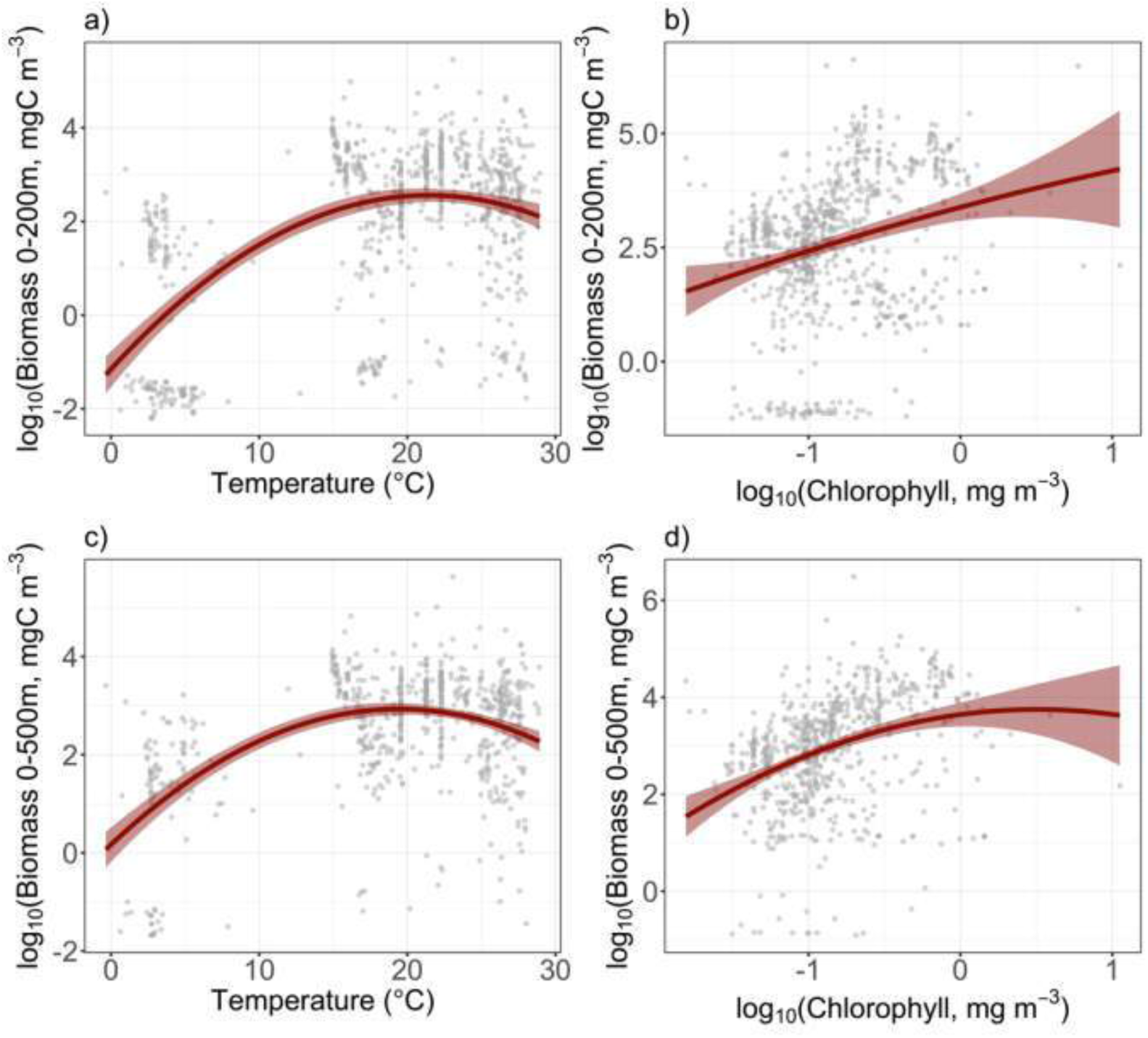
Model fit to each of the predictor variables for giant rhizaria with partial residuals. Top row is 0-200m, bottom row is 0-500m.

##### Total mesozooplankton biomass

For each grid square, we use temperature, bathymetry and chlorophyll and the wet weight factor level to calculate the depth-integrated wet weight biomass of mesozooplankton (excluding rhizaria) over the water column in each grid square to get a total global wet weight biomass is 8.85 Pg (95% confidence interval for this estimate is ×/÷ 7.5-fold). For giant rhizaria, we use temperature and chlorophyll and a 10-1 carbon to wet weight conversion (*34*) to calculate a global rhizaria biomass estimate of 2.2 Pg wet weight (95% confidence interval for this estimate is ×/÷ 8.5-fold). This is close to the estimate of 0.204 Pg carbon weight from (*79*), when converted to wet weight. Rhizaria biomass in the top 200m was 1 Pg wet weight (using a 10-1 wet weight to carbon conversion), which is also close to Biard et al.’s estimate of 0.089 Pg carbon weight (*79*). We combine our global mesozooplankton (excluding giant rhizaria) with our global giant rhizaria biomass estimate to obtain a total mesozooplankton biomass estimate of **10.9 Pg** wet weight biomass, with a 95% confidence interval for this estimate of ×/÷ 7.7-fold.

##### Macrozooplankton

Data derive from 36,268 observations from MAREDAT, Moriarty, 2012 (*83*) https://doi.pangaea.de/10.1594/PANGAEA.777398

After quality controlling the data (e.g., filtering NA observations for biomass, volume filtered, depth sampled), we were left with 23,815 observations We also removed about 8000 observations that recorded zero biomass density because most of the observations (34,938 out of 36,268) were converted from abundance to biomass using conversion equations, but only if the observations recorded species-level information (so as to use median species size). This means that many observations were recorded as zero carbon biomass even though there was non-zero abundance recorded, because species-level information was not retained for that sample, meaning they are not true zero observations.

We used a generalized linear model, with a second order polynomial relationship between log_10_ zooplankton biomass, a linear relationship with log_10_ chlorophyll temperature, and a second order polynomial for log_10_ sample depth (Fig. S8). All predictors are significant, and the R^2^ for this model is 13%, with a very wide spread of residuals. Similar to microzooplankton, macrozooplankton biomass declines at high chlorophyll, but also similar to microzooplankton most of the data comes from areas with less than 1mg m^-3^ of chlorophyll a.

**Fig. S8.**
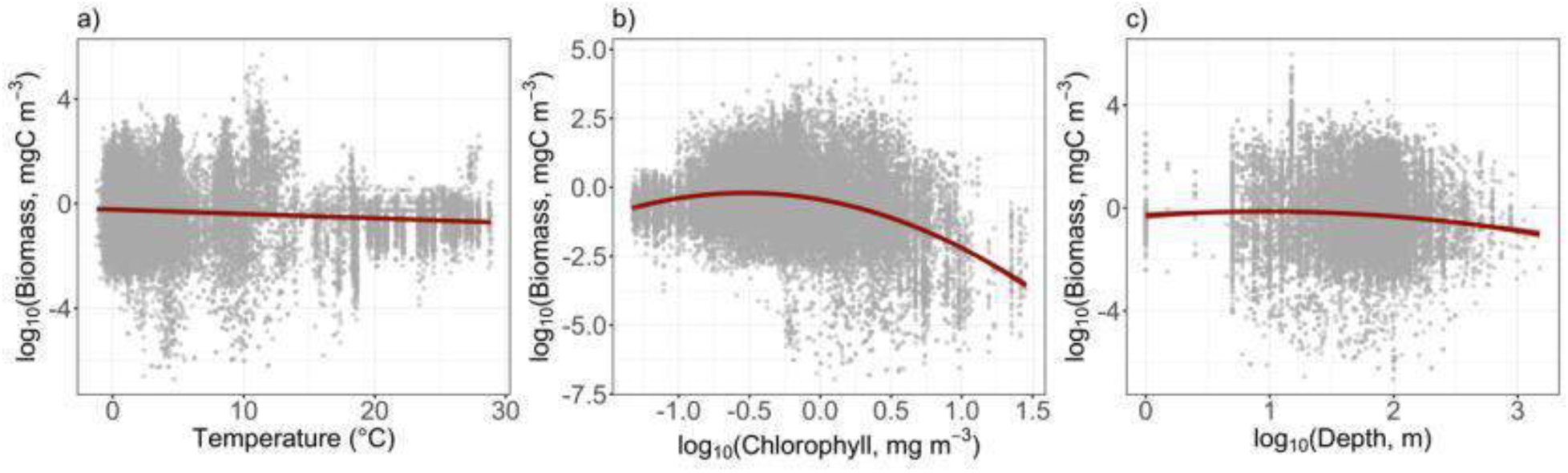
Model fit to each of the predictor variables for macrozooplankton, with partial residuals.

For each grid square, we use temperature, bathymetry and chlorophyll and a carbon to wet weight conversion factor of 10 (*34*) to calculate the depth-integrated carbon biomass of macrozooplankton (g/m^3^) over the water column in each grid square, to get a total global wet weight biomass of **1.5 Pg** (95% confidence interval for this prediction is ×/÷ 11.4-fold).

#### Fish

In the analysis here, ‘fish’ include true fish (bony and cartilaginous) as well as invertebrates of the same size range (mostly cephalopods). Fish include epipelagic, mesopelagic and demersal organisms as well as some benthic organisms that are not covered by zooplankton. We split this group into two groups; mesopelagics and non-mesopelagics, and estimate both separately. We then compare our cumulative estimate of these groups with published estimates to assess how our value compares with other studies.

##### Mesopelagics

Mesopelagic fish live in the mesopelagic zone (200-1000m) of the global ocean. Although difficult to sample directly, these fish can be detected by the reflectance of acoustic signals from sonar, and this reflectance can be used to estimate their biomass (*56*). However, acoustic estimates are sensitive to assumptions about the physiological structure of individual mesopelagic fish, as well as the presence of siphonophores (cnidarians), which have a high acoustic reflectance but low biomass. To address these issues, we used results from (*56*), who developed a model to explore how the percentage contribution of siphonophores to the acoustic backscatter as well as different assumptions about the proportion of fish with gas bladders (which have high reflectance but low biomass), would impact global estimates of mesopelagic fish biomass from acoustic surveys. Taking these uncertainties into account, they reported a range of 1.8 – 16 Pg wet weight (the lower and upper quartile from 500,000 simulations) for the biomass of mesopelagic fish from 70° north and south across the global ocean. Taking the geometric mean of this range, as well as the maximum fold-change from the upper and lower range bound to the geometric mean for our 95% confidence interval estimate, we obtain a global estimate for mesopelagic fish biomass of **5.4 Pg** with a ×/÷ 3-fold 95% confidence interval.

This estimate is similar to (*16*) estimate of 5 Pg wet weight (assuming a wet-weight to carbon conversion of 10), which was based on the geometric mean of a global acoustic survey estimate of 1.5 Pg C (which is probably an overestimate given it assumes most mesopelagic fish have no gas bladder and so have high density) and with a net trawl survey estimate of 0.15 Pg C from (*84*) (which is probably an underestimate given the ability of mesopelagic fish to avoid survey nets; (*50*)). Our estimate is also within the range of uncertainty given by (*6*), who reported a mesopelagic fish biomass between 40° north and south of 2.4 Pg, with approximately a 5-fold uncertainty.

To define the size range of mesopelagic fish, we used the minimum and maximum recorded sizes from three datasets of mesopelagic fish length-weight measurements (*85–87*). The combined datasets contained samples from the trophic and equatorial Atlantic as well as the Southern Ocean and gave a body size range of 0.01-500gm for mesopelagic fish.

##### Non-mesopelagics

For our estimate of the cumulative biomass of epipelagic and demersal fish, cephalopods and large benthic invertebrates, we used spatially-resolved global estimates from two process-based marine ecosystem models (*13, 14*), chosen due to their comprehensiveness and use of data constraints. The first estimate comes from the Bioeconomic mArine Trophic Size-spectrum model (BOATS; (*13*)), a size-based global model that represents harvested organisms (including epipelagic and demersal fish, squid and benthic invertebrates) from 10gm-100kg, using average temperature in the top 75m with integrated primary production in the water column as environmental inputs. The second estimate comes from the FishErIes Size and functional TYpe model (FEISTY; (*14*)), which represents the global biomass of epipelagic and demersal fish, and benthic invertebrates from 1 mg to 125kg, using a functional-type framework, with sea surface temperature, zooplankton carbon concentration and particulate organic carbon flux to the sea floor as environmental inputs. Both models are constrained with catch data, and each are able to reproduce global-scale patterns in catch of pelagic and demersal fish, cephalopods and benthic invertebrates that agree with empirical estimates. These models are members of the FISHeries and marine ecosystem Model Intercomparison Project (FishMIP), and we used publicly available output from these models to derive our global biomass estimates from simulations where each model is forced with environmental inputs from the CESM-BGC1 earth system model (both models’ output available at https://www.isimip.org). For each model, we calculated their global average biomass using the decadal average from 1850-1860 from simulations of pristine ocean biomass with no fishing. We chose this decade because it was the furthest in the past that both models have been run through, and it is far enough back in time that anthropogenic climate impacts are not discernible from existing climate variability (*88*).

Only FEISTY covered the size range from 1mg to 10 g, so we used this model to calculate a global biomass estimate of **1.35 Pg** for pelagic and demersal fish, and benthic invertebrate biomass in this size range. From 10gm to 100kg, the FEISTY model gave a global biomass estimate of 2.25 Pg, and the BOATS model provided an estimate of 5 Pg. Taking the geometric mean of these two values, we estimate the global biomass of pelagic and demersal fish, benthic invertebrates and cephalopods from 10gm to 100kg to be **3.6 Pg**. To obtain the biomass of fish with body sizes greater than 100kg we used the scaling relationship between biomass and body size from (*29*), who used macroecological theory from (*89*) to define the relationship between biomass and body size as a power function with an average exponent of −0.06 (assuming an average ecosystem trophic transfer efficiency of 0.125, and an average predator-prey mass ratio of 1000). According to this scaling, total biomass in any given order of magnitude size range is 88% of the biomass in the preceding order of magnitude range. Using this relationship, and their reported median biomass of all fish between 1gm and 1 tonne of 4.9 Pg, we were able to obtain a global biomass estimate of total biomass from 100kg-1 tonne (covering groups such as adult salmon and tuna) of **0.57 Pg**, from 1 tonne-10 tonne (covering groups such as sunfish, rays and sharks) of **0.48 P**g. The fish component of this biomass was then calculated by subtracting mammal biomass (derived below).

To calculate the uncertainty range for our estimates of non mesopelagic fish biomass, we follow the approach of (*16*), who took the 90% confidence interval of global fish biomass from (*90*) as representative of the uncertainty of their non mesopelagic fish biomass estimate. Jennings and Collingridge (*90*) calculated a median biomass of fish between 1e-5g and 1 tonne of 14 Pg, with a 90% confidence interval of 2.9 Pg to 46 Pg. Using this interval, we calculated a ×/÷ 5-fold 95% confidence interval for our estimate of non-mesopelagic fish.

##### Comparison of total fish biomass with other global estimates

When combined, our estimate of global fish biomass is within the range of estimates reported in previous studies. For example, our estimate of total biomass from 1g to 1 tonne is 8.2 Pg, almost double the 4.9 Pg estimate of (*90*) but within the 50% uncertainty interval for their estimate (2 – 10.4 Pg). More broadly, for all organisms from 1e-5g to 1 tonne, covering mesozooplankton, macrozooplankton and all fish, we obtain a global biomass estimate of approximately 23 Pg, which is almost double the 14 Pg estimate of (*90*), but still within the 50% uncertainty interval of their estimate for this size range (7.7 Pg – 23 Pg). Finally, (*16*) estimated the global biomass of mesopelagic and pelagic fish, benthic organisms and cephalopods to be approximately 10 Pg wet weight (assuming 10-1 wet weight to carbon conversion), which is close to our total estimate for these groups of 10.9 Pg.

**Table S5.**
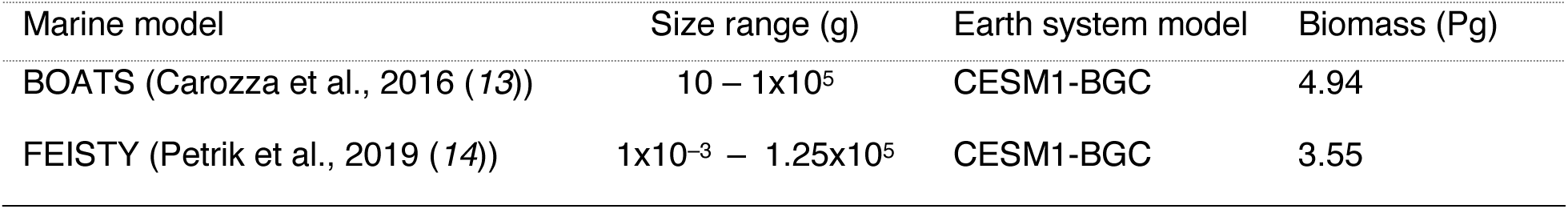
Summary of fisheries models, their size ranges and global fish biomass estimates.

| Marine model | Size range (g) | Earth system model | Biomass (Pg) |
| --- | --- | --- | --- |
| BOATS (Carozza et al., 2016 (13)) | 10 – 1x10<sup>5</sup> | CESM1-BGC | 4.94 |
| FEISTY (Petrik et al., 2019 (14)) | 1x10<sup>-3</sup> – 1.25x10<sup>5</sup> | CESM1-BGC | 3.55 |

#### Mammals

Marine mammal biomass was estimated from species-specific surveys for the majority of all known species. These are usually underestimates due to limited observation over portions of a species full geographic distribution, or else due to the inherent difficulties of counting individuals that dive for long periods or actively evade observers. Moreover, we seek both current and “pristine” numbers, the latter of which might characterize the pre-whaling period, before any observations of population numbers were systematically taken. This necessarily involves guesswork, but the range of possibilities is narrowed by making use of multiple independent estimates for the current abundance, body mass, and geographic range, along with growth rates and human capture data. These complementary data are available for most marine species, as well as for most non-marine mammals, against which the marine estimates can be compared.

Population data of varying quality exist for more than 2/3 of all ∼ 126 extant species of marine mammals. Nearly all current estimates of marine mammals are from IUCN (*57*) (*n*=87). We estimated “pre-whaling” population counts by taking larger valued estimates or upper confidence intervals when extensive capture levels have been reported, or else from expert opinion or modeling studies (*n*=83). For many of the larger whales, we relied on estimates in (*91*), which is in part based on a study by (*28*). When no additional information was available beyond current estimates, we assumed pristine numbers were the same as exist currently. We believe this approach is more likely to underestimate abundance than overestimate it in most instances, but they provide among the most complete tallies of currently available data for marine mammal biomass yet assembled.

To provide population estimates for the 1/3 of species that have not been surveyed, we used regression predictions for the 2/3 of species for which observations are available. These data and associated species body mass and geographic range area were used to build relations between mammal population density and body mass. By dividing global population counts by geographic range area, we obtain mean population density, which we regressed against body mass for the 2/3 of marine mammals for which empirical counts exist (Fig. S9 A). These regressions were used to estimate pristine and current densities for the remaining 1/3 of species for which no current or prior estimates exist. Mean body mass estimates were obtained from (*92*), while range areas were obtained from shapefiles from (*57*) and (*93*) for all 126 marine mammal species. This allowed us to estimate the population density over their range for the remaining 40 some species (yellow points in Fig. S9 A), and to calculate their total numbers by multiplying by geographic range. While marine mammal species generally exhibit poor relations between global count vs. body mass (Fig. S9 B), geographic range area vs. body mass (Fig. S9 C) and global count vs. geographic range area (Fig. S9 D), their population density vs. body mass relation (Fig. S9 A) is comparatively robust for both the pre-whaling period and current estimates. Nonetheless, these regression predictions for species abundance without extensive surveys tend to be considerably higher, sometimes by orders of magnitude, than the estimates listed in (*28*), most of which are species with the lowest confidence and limited survey over their geographic distribution (see Supplementary Data file).

**Fig. S9.**
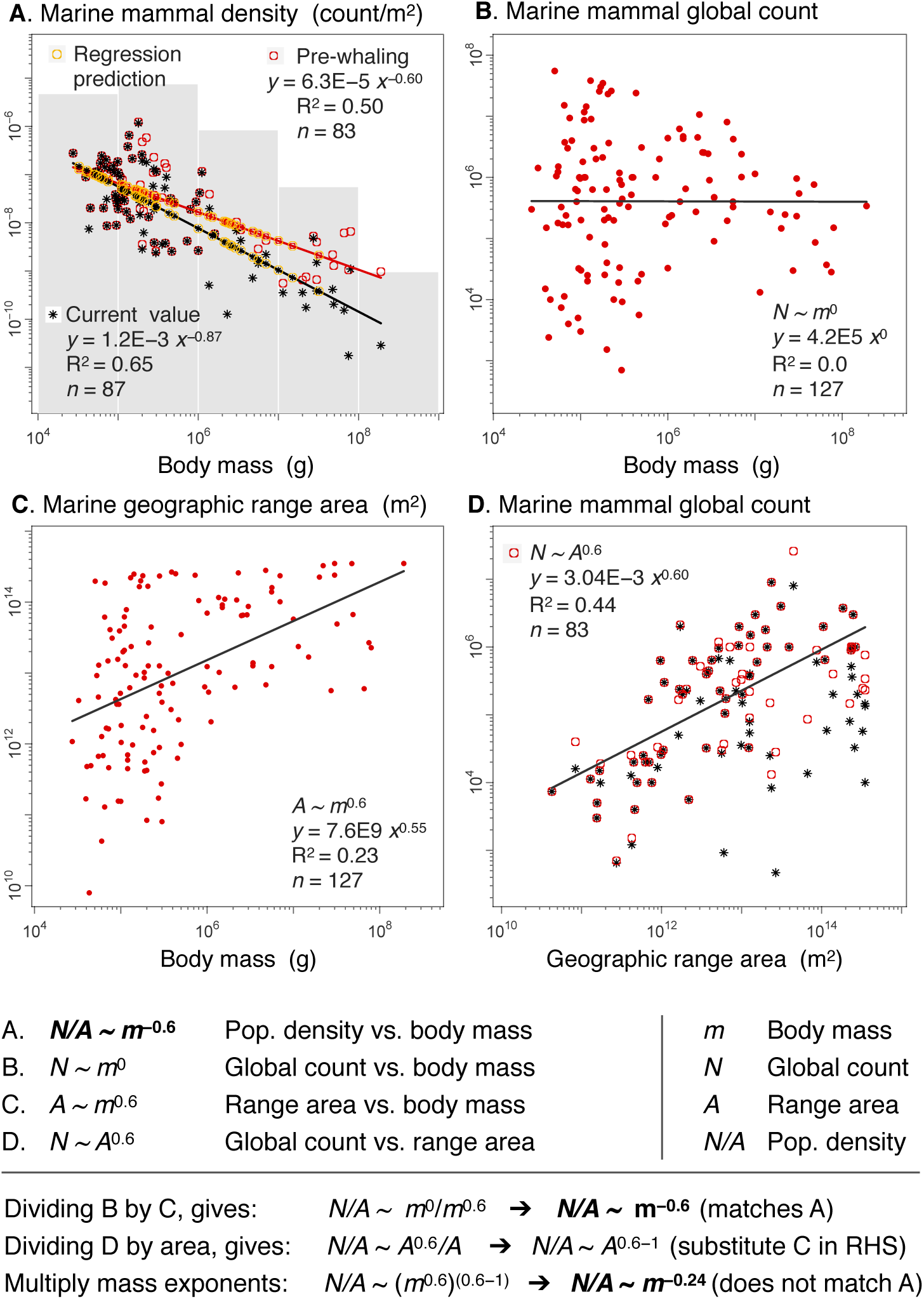
Marine mammal population regressions. **A.** Mammal population density is calculated by dividing the global species count by geographic range area. Regressions for both pre-whaling estimates (red circles) and current densities from IUCN (black stars) allow species with no global data to be predicted (yellow circles). The grey bins sum total marine mammal population density in each size class. **B.** Total global counts of mammal species are uncorrelated with body mass (*N ∼ m*^0^). **C**. Mammal geographic range area shows a positive relation with body mass (*A ∼ m*^0.6^). **D**. Global mammal population counts show a positive relation with geographic range area (*N ∼ A*^0.6^). Regression line is to pre-whaling values (red circles) and we did not account for shifts in range size. Regressions B to D show considerable scatter, giving different predictions for the regression in A. For example, pre-whaling density (A; *N/A ∼ m*^−0.6^) is consistent with combining B and C, (*N/A ∼ m*^−0.6^), but is not consistent with combining C and D (*N/A ∼ m*^−0.24^).

Mammal population densities and population biomass are also not notably different from terrestrial species, but tend to fall below the terrestrial body mass regression for both pre-whaling and particularly for current estimates (Fig. S10 A and B). The regression of terrestrial and pre-whaling densities has an exponent of −0.9, which is considerably steeper than what has been shown over a more limited range to be near −0.75 (*10, 94, 95*). Marine mammals, and particularly whales, extend the terrestrial mammal size-density relationship by a full two orders of magnitude.

**Fig. S10.**
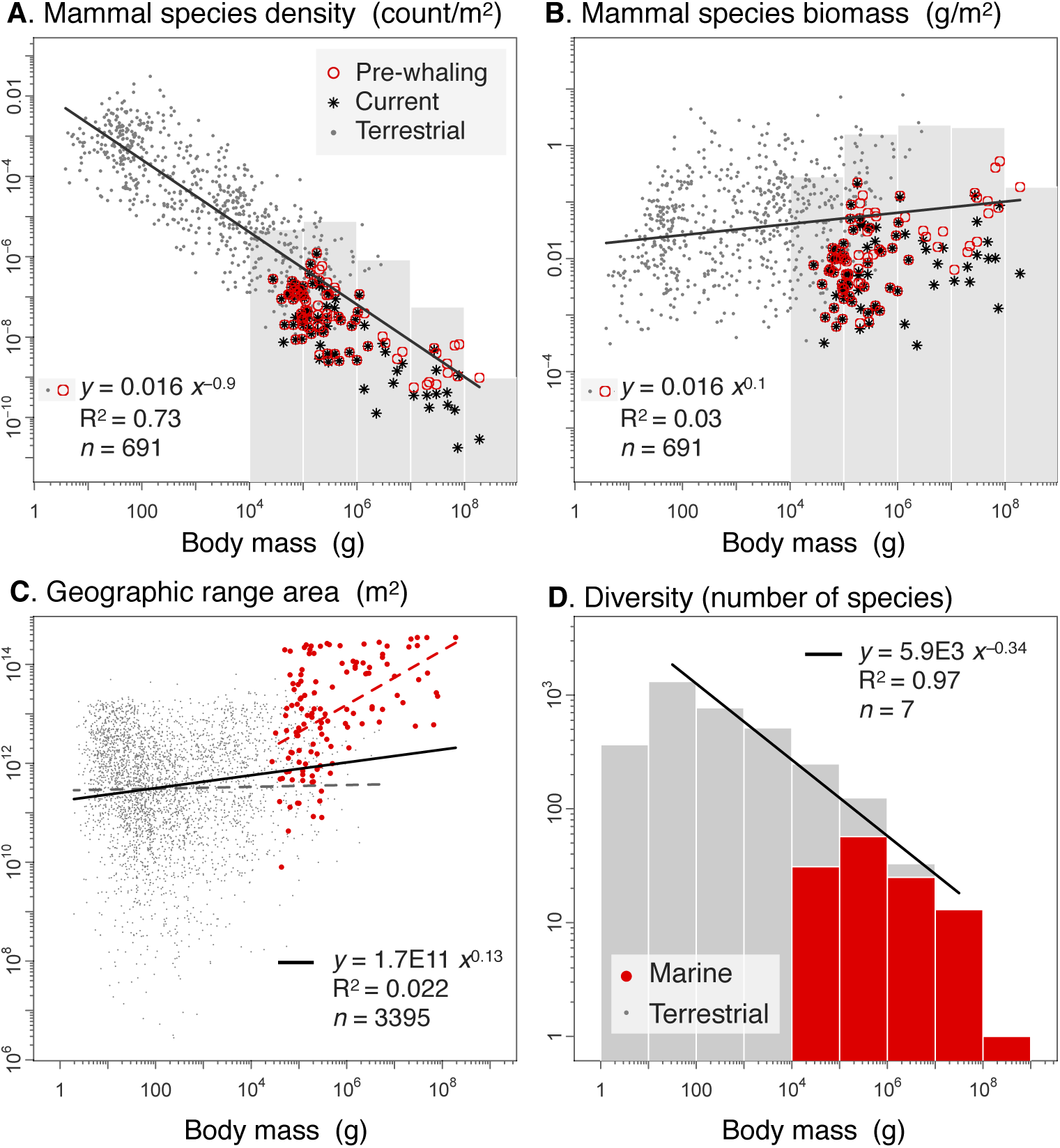
Comparing marine and terrestrial mammal body mass regressions. **A.** Marine mammal population density from Fig. S9 A are compared with (mostly) terrestrial mammal species (grey circle). The regression line is for terrestrial and pre-whaling estimates (excludes current densities). **B**. Mammal population biomass is calculated by multiplying densities from A and body mass. The grey bins represent total marine mammal population density sums over each order of magnitude size class. **C.** Marine (red, from Fig. S9 C) and terrestrial (grey) mammal global geographic range areas. **D.** Mammal diversity (species richness) across order of magnitude size classes. The regression line excludes the smallest and largest (blue whale) size classes.

Marine mammal abundances may be underestimated for many reasons, including i) difficulties in counting individuals hidden deep under the surface, ii) poor estimates of geographic range areas used to estimate densities or iii) centuries of human capture. Disentangling these many factors is not possible with currently available information. For reference, we also show how geographic range area vs. body mass compares between terrestrial and marine mammals, with all body masses capable of occupying the entire surface of land or sea, but larger animals apparently incapable of occupying very small areas (Fig. S10 C). We also show for all mammal species how species diversity (count of species in each size class) compares between terrestrial and marine species (Fig. S10 D).

Our measure of uncertainty for large mammals derives from considering extensive additional data that we include in our dataset, but are not shown in the figures. In addition to “pre-whaling” and current best estimates of marine mammals, we made use of additional current estimates of marine mammals also listed on IUCN species pages (*57*), as well as data drawn from Appendix 4 of (*28*), which is derived from some 275 published sources, and lists minimum, mean and maximum values for 115 of 126 marine mammal species. Many of these estimates are reportedly of very low confidence over a limited geographic range, and differ from our current best estimates to varying degrees, but tend to be within a factor of three of current values. It is more difficult to obtain a measure of uncertainty for pre-whaling abundance, but even the most extreme species specific values are rarely more than a factor of ten different from other estimates, and on average are within a multiplicative factor of 4.3 of maximum current values from (*28*). Pooling all estimates including those of (*57*) and the minimum, mean and maximum current values of (*28*) gives a mean fold uncertainty of 2.2 (following (*15, 16*)). However, these separate estimates are not necessarily independent, and our knowledge of the pre-whaling period is necessarily vague. We considered a fold uncertainty value of 5 to be cautious, and include all additional marine mammal estimates in the Supplementary Data file.

Further assumptions about how we chose to deal with uncertainty and our regression methods are covered in the section “Statistical considerations”, below.

### Human impacts on the biomass spectrum

The most dramatic human impacts on marine biomass to date have been the hunting of wild animals, including fishing, sealing, and whaling. Climate change, on the other hand, is rapidly growing in importance and is expected to be a major driver of change in abundance over the current century. Because dramatic fishing impacts have already occurred and regulation efforts are now slowing the rate of overfishing in many parts of the world (*96*), we use empirically-informed estimates of the impact that fishing has had to date, focusing on the depletion caused by industrial fishing up until the early 2000s. We compare this with the climate change impacts expected under a pessimistic scenario for the 21^st^ century, assuming that the impacts of fishing and climate on biomass are approximately additive, with little non-linear interaction between them (*97, 98*). Thus the estimated historical fishing impacts are summed with projected climate changes to provide an estimate of the total impacts that would occur in future if the effective fishing effort remains approximately constant. Although humans have also impacted biomass by physically modifying habitats, most importantly coastal regions, estuaries and waterways used in anadromous fish migrations (*99, 100*), we do not address these directly.

In Fig. 3 A, we present human biomass values obtained from the literature. Annual mammal and wild fish harvests were estimated at 0.12 Gt from (*51*), while total annual human consumption is estimated at 4.7 Gt from (*101*), making human harvests over the ocean a small fraction < 3% of what humans consume globally. We use values for total human population of 1 billion in 1800, 8 billion currently and 11 billion in 2100. We multiply these values by a reference human being of 60 kg to get biomass values for humans shown in Fig. 3 A.

#### Fishing impacts

Fishing has had a major impact on the biomasses of fish (including true fish as well as targeted invertebrates). The available estimates that have been made for the global biomass depletion by fishing do not explicitly address changes in the distribution of biomass depletion over the size-spectrum, however, some local studies have addressed the size-distributed impact. Thus, we first consider the global fish biomass depletion, and second the changes in the size distribution based on local observations.

##### Biomass decline

There are many local and taxon-specific estimates of the decline of fish biomass caused by fishing activity, but relatively few global estimates. Furthermore, the long history of modification of ecosystems by fishing, which often began long before data was collected, leads to a problem of shifting baselines (*102*), making it difficult to reconstruct the pristine state. We include two different global estimates that approximately target the timespan of 1950 to 2000, each of which is methodologically independent.

1. Worm and Branch (*103*) present a compilation of fish stocks assessed by scientific management agencies. These covered only 20-25% of fished stocks at that time, but suggested that, for these stocks, only 31% of the pristine biomass remains. This is consistent with fishing pressure being exerted slightly in excess of that which would produce maximum sustainable yield, generally expected at roughly 40% of pristine biomass (*104*). Fishing pressure generally occurs on fish >10 g, and so this estimate is applied across all fish larger than 10 g. A similar value is obtained from (*105*) using a process model of the global fishery, including all potentially commercial fish >10g. These estimates are subject to uncertainty in the unfished biomass fraction, i.e. the proportion of the total fish biomass that is not harvested. This unfished biomass is not included in stock assessments, and is absent from global fishery models. We are not aware of published comprehensive assessments of this unfished fraction, but a recent model-based study provides a global estimate that 58% (±22%) of fish biomass is targeted by commercial fisheries (*106*), and that 42% is not targeted. In the absence of empirical data, we use this model estimate of commercial fisheries fraction to calculate the total fractional decline in fish biomass as follows: the commercial fisheries decline (0.31) multiplied by the commercial to total fisheries fraction (0.58), plus the fraction of fish not targeted by fishing (0.42), gives an estimated 60% of total fish biomass >10g remaining from pristine levels.

2. Christensen et al. (*107*) provide a compilation of more than 200 Ecopath models constructed for globally-distributed ecosystems and incorporating data on fish biomass. They fit a multi-variate linear regression to the simulated biomasses of predatory and forage fish in order to estimate the changes over time. They estimate a decrease in global biomass of all predatory fishes by two-thirds (60.2 - 71.2 %, 95% C.I.) over the century from 1910-2010, thus leaving only 33% of fish biomass >10 g remaining from estimated pristine levels. Christensen et al. (*107*) do not provide a size estimate for these fish, but here we assume this applies to all fish >10 g, commercial and non-commercial. They also suggest that small forage fish have doubled in biomass, potentially by as much as 130%, though we have not incorporated this suggestion, which remains an intriguing possibility.

We combined these two estimates for the fraction of fish >10g remaining from pristine fish biomass (60% inferred from (*103*), and 33% inferred from (*107*)) by taking their geometric mean, giving a value of 45% of fish >10g remaining.

##### Impacts on the size structure

As first demonstrated by (*108*), fishing has very strong effects on the size structure of exploited species and associated bycatch for three main reasons, all of which cause a steepening of the size-spectrum. First, larger fish are preferentially targeted given that most gears select sizes larger than some mesh cutoff size, increasing bycatch of larger organisms, and larger fish tend to be more desirable (*109*). Second, because large fish take many years to grow slowly from smaller fish, high rates of exploitation of young fish prevent the replacement of large, older fish (*13*). Third, because heavy exploitation favours quickly-reproducing individuals, there are strong evolutionary pressures towards smaller maximum sizes (*110*). Jennings and Blanchard (*110*) show trawling data for the heavily exploited North Sea, revealing a biomass slope of −1 (abundance slope of −2) over the range of 1-66 kg, which they interpret as a sharp steepening of the pristine slope of this size range of the benthic community by −1. Together, these factors have been shown to produce strong increases in the slopes of size spectra in most exploited communities (*2*).

We therefore calculated what size-spectrum slope would be consistent with the 45% depletion among the fraction >10g, assuming that fish ≤10g have not been significantly affected by current harvesting in the global average. This would require that the slope has become steeper by −0.17 for all fish >10g. Thus, we assumed no change in the size-spectrum for fish ≤10g, but above this the size spectra slope has been steepened by −0.17, account for the inferred fraction of fish remaining relative to pristine levels.

We note that different ecosystems seem to show different sensitivities of the size-spectrum to fishing. Demersal fish, which may have steeper size spectra slopes, appear to be relatively sensitive to harvesting (*111*). In contrast, coral reef size-spectrum slopes tend to be less steep, with baseline slopes > −1 and proving relatively robust to fishing pressure (*112*). Thus it appears likely that coral reef size structures have been less impacted than the global average, possibly due to the stabilizing effect of very large grazers. These observations serve to emphasize that our global inferences for fish declines will not likely apply to particular ecosystems and are necessarily a first order approximation.

In addition to steepening in the size-spectrum due to fishing, there have also been significant declines in large mammals, particularly whales. As described in the section “Data and methods” under “Mammals”, along with accompanying data at the github link provided in the first paragraph of this file, we compiled both pre-whaling and current abundance for more than 2/3 of all ∼ 126 extant species of marine mammals. These data derive from over 275 primary sources, as well as meta-analyses of (*28, 91*), as well as expert opinion and compilation from (*57*). While these data do not include all species, and are more limited for pristine species biomass estimates, they summarize the state of current knowledge and suggest some startling declines particularly among the largest size classes. In particular, these data suggest that among the smallest marine mammals (10-100kg), only 47% remain, but that progressively smaller fractions characterize the larger size classes, with possibly only 25% of mammals remaining in the 10 to 100 tonne size class, and only 3% of blue whales, making up the largest size class (*91*).

Combining the declines due to fishing and whaling results in significant changes of the size-spectrum, not only among harvested size classes, but with noticeable effects on the overall slope from bacteria to whales, as shown in Fig. 3 C and Table S6 (compare i and ii).

**Table S6.**
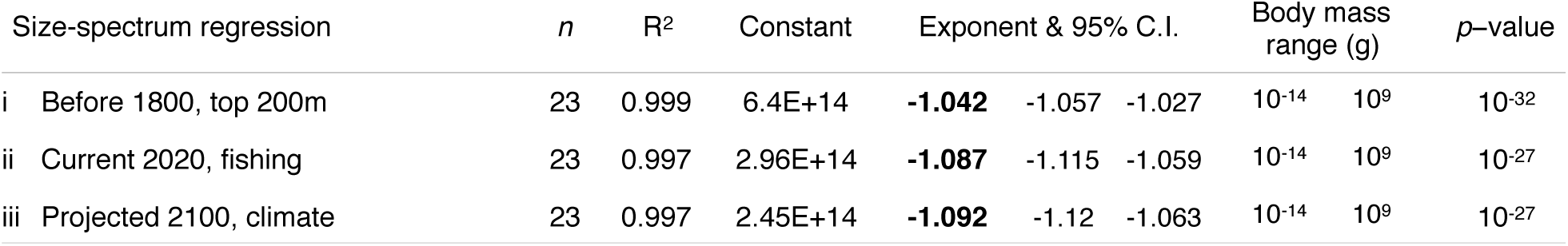
Human impacts on the size-spectrum. The global ocean regression (i) represents the estimated pristine (before 1800) size-spectrum, while regression (ii) is the size-spectrum estimated currently (2020) that includes the impacts of fishing, and (iii) is estimated under climate projected impacts to 2100.

#### Climate projection impacts

To provide an illustration of expected impacts of climate change on the full size-spectrum of marine life, we assembled estimates of relative changes in biomass obtained by previously-published model projections for different taxonomic groupings. In all cases these studies compared the last decade of the 21st century with the last decade of the 20th century – thus, these do not estimate the total impact of climate change from the pre-industrial state, but rather serve as a consistent metric in the literature that is used here to gauge relative impacts across size.

At the root of all these projected changes are Earth System model simulations, that estimate the changes in ocean temperature, circulation patterns and biogeochemical cycling that will result from a given future trajectory of atmospheric greenhouse gases. Here we use the worst-case scenario trajectory, Representative Concentration Pathway (RCP) 8.5.

Rather than rely on a single model, we take the mean estimate from the ten climate models that participated in the fifth simulation round of the Climate Model Intercomparison Project (CMIP5) and included biogeochemical components, as analyzed by (*113*). The multi-model mean sea surface temperature (SST) changes from 1990-2000 to 2090-2100 under RCP8.5 is 2.73 ±0.72 °C. This temperature change would be expected to be representative of the average warming within the continually-mixed surface layer (the upper 20-80 m). At greater depths the warming would be less, given the time required for heat to penetrate from the atmosphere to the subsurface.

Changes in plankton concentration were calculated by the same Earth System models that estimate the physical changes, using ecosystem modules that were run in a fully-coupled mode with the ocean modules. According to the analysis of (*114*), these models simulate a global mean phytoplankton decrease of 6.1% ± 2.5% and a global mean zooplankton biomass decline of 13.6 ±3.0 %. These changes were applied uniformly across the full size ranges of phytoplankton and zooplankton, respectively.

Climate-driven changes in fish biomass are also provided by global dynamical process models, run offline using the output of the Earth System models. Typically the most important drivers are net primary production and water temperature. The multi-model analysis of (*98*) projected a 17 ±10.7% decrease in animal biomass, for the resolved animals >10 cm.

Unlike the case for the preceding groups, there are no global process models available for marine mammals. Instead, we use the statistical model of (*115*). These authors argued for a competitive niche of marine mammals that shrinks with rising temperatures, so that a 1 °C increase of sea surface temperature results in a 12% decline of marine mammal abundance (24% for pinnipeds). Thus, the CMIP5 estimated warming of 2.73 °C implies a 30% reduction in marine mammals and a 53% reduction in pinnipeds.

Finally, we were unable to find prior estimates for the relative changes in bacteria concentrations under climate change. Instead we estimated the changes using the temperature and chlorophyll dependence from the statistical model we developed to estimate global bacteria abundance, an approach analogous to that used by (*115*) for marine mammals. We obtained decadal averages of global sea-surface temperature and surface chlorophyll-a concentration for 1950-1960 and 2090-2100 from the GFDL-ESM2M and IPSL-CM5A earth-system models, forced with historical and RCP85 scenarios. We used these decadal averages as inputs to our statistical model of global bacteria abundance, and obtained an estimate of the relative change in bacteria abundance in 2090-2100 compared to 1950-1960 under the RCP85 warming scenario for each earth-system model. Contrary to other major groups, we estimate that global bacteria abundance will see a moderate increase of approximately 2.3% by the end of this century under an RCP8.5 warming scenario, compared to 1950-1960 levels.

While climate change projections could have significant unanticipated changes on marine ecosystem function, they appear to pale in comparison to the impacts on the size-spectrum resulting from fish and mammal harvesting, as shown in Fig. 3 B and Table S6 (compare ii and iii).

### Statistical considerations

In this section, we outline the various ways of representing the size-spectrum, our fitting methods, and the sensitivity of our assumptions of group size distributions and group biomass to our overall findings.

#### Fitting size-frequency distributions

Unlike the size-frequency distribution of people, which follows the classic ‘bell-curve’ or normal distribution, a power law distribution is extremely right skewed, also referred to as heavy-tailed or fat-tailed. The power law distribution (also called the Pareto distribution) is more like a log-normal distribution but with no central mode, such that the smallest size class is overwhelmingly the most frequent, and the largest size class is extremely rare, but orders of magnitude larger than the smallest (*116–118*).

A power law is revealed as a straight line when observations are plotted on log-log axes of size and frequency. The same data appears as a capital “L” on regular axes, and is not informative over larger size ranges. Many claims of underlying power law mechanisms have been made from the appearance of a straight line on log transformed axes, some of which are undoubtedly false (*119*– *122*). In the past decade, methods have been developed based on maximum likelihood that better evaluate if the underlying data conform to a power law based on more principled statistical methods, and claim to estimate the exponent with greater precision than ordinary least squares. In addition, these methods make use of Kolmogorov-Smirnov (KS) goodness-of-fit tests and Vuong’s likelihood ratio tests with other long tailed distributions (*119*). Unfortunately, there has been considerable criticism of these methods (*120–122*), implying that the statistics of fitting power law distributions has failed to reach broad consensus.

An additional source of confusion is the proliferation of different ways of constructing these size-frequency distributions, which is not only extensive among aquatic ecologists, but also among the many other fields of science that study power law frequency distributions. Here we list several of the many ways that the size-spectrum can be represented:

*- Size-spectrum or abundance-spectrum*: This is the size-frequency distribution shown in Fig. 1, representing numerical abundance (y-axis) vs. size class (x-axis), both in logarithmic scale (including logarithmic bin width). This is a common way that the distribution of individual sizes is represented. A size-spectrum slope of −1 indicates that size and abundance are inversely proportional, i.e. individuals decrease in number by the same factor as they increase in body mass.

*- Biomass-spectrum*: This is the size or abundance spectrum multiplied by the body mass size-class, as shown in Fig. 2, and is also a common way that these distributions are represented. For example, a biomass-spectrum slope of 0 is equivalent to an abundance-spectrum slope of −1. Biomass may be represented on logarithmic or linear axes, depending on the study.

*- Normalized size-spectrum*: This is the size-spectrum but the count of individuals in each bin is divided (or “normalized”) by the bin width (although bins appear to have equal width on log axes, they are in fact getting multiplicatively larger as size increases) (*2*). A normalized size-spectrum slope of −2 is equivalent to an abundance-spectrum (with log bins) slope of −1. The normalized size-spectrum exponent is equivalent to the probability density function exponent, though the former is a discrete distribution while the latter is continuous (*118, 123*).

*- Normalized biomass-spectrum*: Similarly, this is equivalent to the biomass spectrum except that the biomass in each size-class is normalized (divided) by bin width. A normalized biomass-spectrum slope of −1 is equivalent to a biomass-spectrum (with log bins) slope of 0.

*- Cumulative distribution*: Instead of binning the data, the distribution can be expressed as the probability that a randomly selected individual is greater than or equal to a given size. The cumulative distribution is sometimes thought to be a superior way of representing the data, given that it eliminates the requirement for arbitrary decisions about how to construct bins (*118, 119*). However, for some datasets a cumulative distribution can appear very smooth, while a histogram may better reveal variation between size classes.

*- Rank-frequency distribution*: Instead of plotting the cumulative distribution on the y-axis and size on the x-axis, we can instead plot the rank of the cumulative distribution against body mass (so that it would be more accurately called a rank-size plot). This is typically done by reversing the axes, so that rank is on the x-axis and size is on the y-axis. This is also known as Zipf’s law (*117, 118, 124*). If we rank the sizes of individuals in order, then there are *n* individuals with frequency greater than or equal to that of the *n*th largest size. The rank and cumulative distribution are thus proportional, and since the axes have been reversed then the exponents of the distributions are inverse, which in the particular case of a slope of −1 does not matter (see also (*125*).

In addition to these different ways of expressing the size distribution, ecologists have expressed size on the x-axis in terms of length (e.g. diameter), volume or mass (e.g. wet or dry mass) or energy density (e.g. kilocalories or watts). Although the exponent is unlikely to be much affected by conversions between volume, mass or energy density, the use of length as a measure of size yields very different exponent values. For example, if the size-spectrum uses length, and has a cumulative distribution function slope of −α, then to convert to volume, the slope will be −α/3. If the probability density function uses length and has a slope of −α, then to convert to volume, the slope will be –(α+2)/3. Although Sheldon et al. originally represented size in terms of length (equivalent spherical diameter; Sheldon et al. 1972, Sheldon and Parsons 1969), they binned their data in units of volume, so that converting length to volume would not alter the exponent. It is not clear, however, that the same kind of volume based binning has been applied in other studies leading to significant confusion.

While alternative fitting methods and representations of the size-spectrum are important to be aware of, several are less relevant to our purposes. The earliest plankton size-spectra, and the many that came after, were constructed from passing a water sample through a Coulter counter, which measures the size of every single entity over a respectable six orders of magnitude mass range, with current coulter counters extending up to possibly ten. The data are thus in the form of a very long list of different sized individuals that can be then binned in various ways, or ranked or fashioned into a cumulative distribution, the latter of which is often the preferred form for approaches based on maximum likelihood (*119–122*). In contrast, our global estimates are binned, with the allocation of different groups to bins themselves having some uncertainty bounds. Indeed, we cannot know the sizes of all the individuals in the global ocean, a list of which would be on the order of 10^30^ (or ∼2^100^) elements long.

Despite these limitations, our data extends over a size range of some 23 orders of magnitude. Very few power laws in other fields of science, with the exception of perhaps astronomy, come anywhere close to these size ranges (e.g. see list in (*118*)). Many claimed empirical power laws that have later been claimed to be spurious under alternative statistical approaches typically span only two to five orders of magnitude. Our purpose was to show that the Sheldon hypothesis is largely validated (with two broad exceptions being bacteria and whales), rather than to provide precise estimates of the overall slope. Nonetheless, our best estimates of abundance in each size class exhibit a highly regular pattern over this extraordinary size range, such that different estimators of the exponent are quite similar, as shown in Table S7.

**Table S7.** Comparing exponents from different fitting methods. Regression statistics for the global ocean and that restricted to the top 200m are shown for ordinary least squares (OLS), with exponents from maximum likelihood (MLE) and reduced major axis (RMA) shown for comparison. “*n*” refers to the number of size class bins, R^2^ is the coefficient of determination, “Constant” is the abundance when size is 1 g, and 95% C.I. is the 95% confidence interval for the OLS derived exponent.

| Size-spectrum regression | *n* | $R^2$ | Constant | Exponent | | | 95% C.I.<br>(OLS) | Body mass<br>range (g) | | *p*-value |
| --- | --- | --- | --- | --- | --- | --- | --- | --- | --- | --- |
|  |  |  |  | MLE | RMA | OLS |  |  |  |  |
| ii Global ocean, top 200m | 23 | 0.999 | 6.61E+14 | **-1.028** | **-1.040** | **-1.040** | -1.054 -1.026 | 10<sup>-14</sup> | 10<sup>9</sup> | 10<sup>-32</sup> |
| ii Global ocean to seafloor | 23 | 0.998 | 1.09E+15 | **-1.033** | **-1.052** | **-1.051** | -1.072 -1.03 | 10<sup>-14</sup> | 10<sup>9</sup> | 10<sup>-27</sup> |

We show statistics obtained from ordinary least squares (OLS), and compare these exponents with those of maximum likelihood (MLE; following methods outlined in (*121*) for binned data), and reduced major axis (RMA; following methods outlined in (*126, 127*)). The latter method partitions error in the data equally to both x-and y-axes (unlike OLS, which assigns all error to the y-axis), and so is not really appropriate for our data. Although there is undoubtedly some error in our assignment of different major groups to different size classes, the vast majority of our uncertainty lies in our ability to estimate abundance over the global ocean in each size class. Nonetheless, we report RMA exponents for reference, and note the similarity in the exponents among all methods (OLS, MLE and RMA).

#### Sensitivity analysis of group size distributions

Most research over the past 50 years has revealed that the biomass distribution within major groups is approximately invariant with body mass (Table S3, and references therein). Nonetheless, to make such an assumption *a priori* across each of our 11 major groups (see Table S2) would potentially weaken our ability to test the hypothesis of invariant biomass across all groups together. As we show, however, alternative assumptions about the size distribution within major groups, even those that are quite unrealistic, all give very similar global patterns from bacteria to whales.

For example, instead of assuming an invariant biomass within each major group, we can assume a log-normal distribution of biomass across each group, for which there is little evidence of which we are aware, but could nonetheless serve as a valid null hypothesis. We assumed the size range of each major group corresponds to four standard deviations, and that the total biomass independently estimated for each group constrains the distribution. We also binned groups into half order of magnitude width, though this has a negligible effect on our results. We show that the slope for the size-spectrum and the biomass spectrum is essentially unchanged from Figs. 1 and 2 (see Fig. S11). Similar analysis using other long-tailed distributions and binning criteria yield little differences in slope across all groups.

**Fig. S11.**
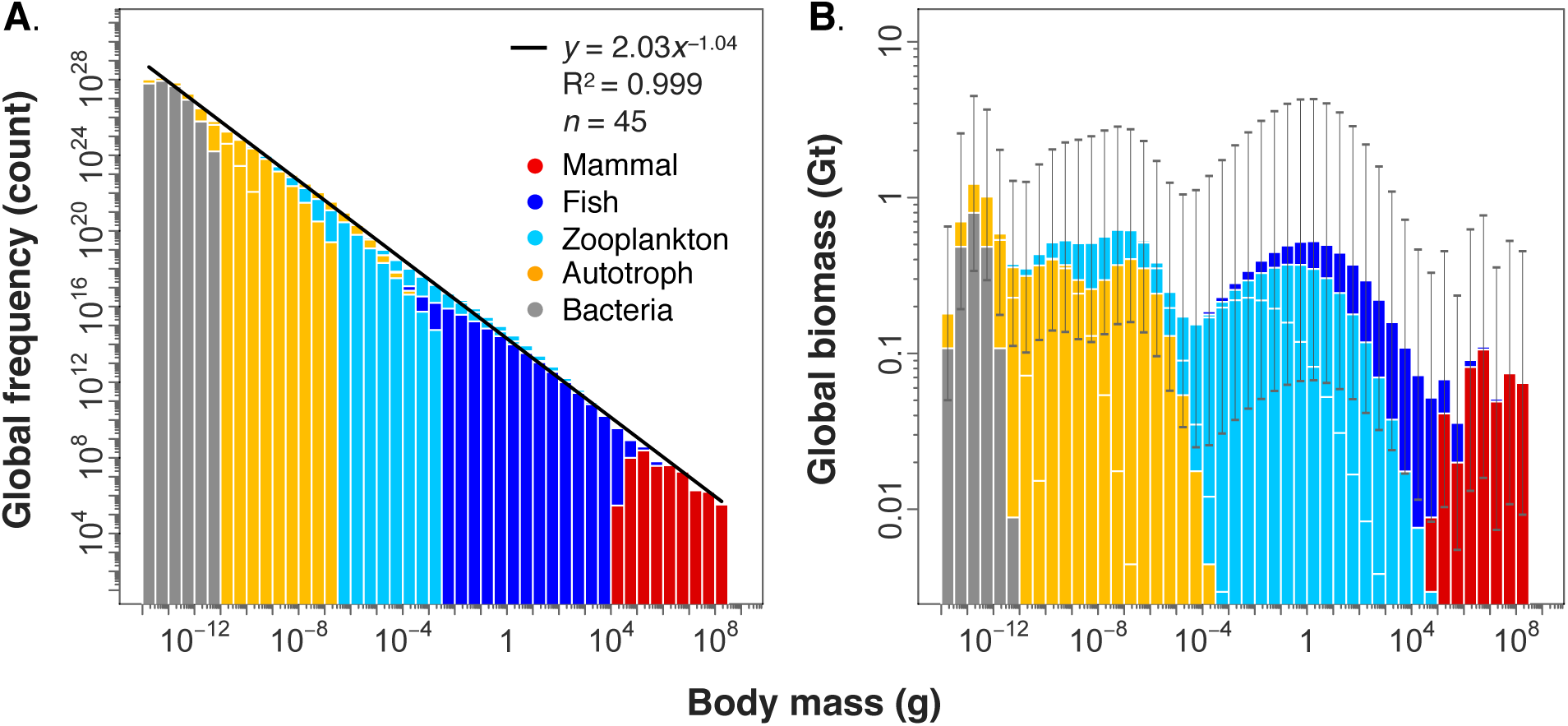
The size-spectrum assuming a log-normal distribution of biomass. Total global frequency (**A**) and biomass (**B**) vs. body mass size-class assuming each of eight major groups are log-normally distributed with respect to size. We have assumed that our best estimate of the logarithm of size range for each group corresponds to four standard deviations. The bin width can be modified arbitrarily without altering the slope (shown here are half an order of magnitude size bins). Depending on how groups are stacked within a given size class determines which group is given prominence on the logarithmic plot (plots in A and B are the same data, but are stacked differently so that they appear different).

We have undertaken sensitivity analysis on the size distribution of major groups and find that even quite extreme variation in group size distributions still yields slopes centered on the slopes reported in Figs. 1 and 2, with little variation about the mean.

When we assume that the biomass in each group is uniformly distributed within each group’s size range, we obtain a slope of −0.060 for the total depth integrated global biomass size-spectrum, and −0.040 for the global biomass size-spectrum in the top 200 m. To assess the uncertainty of these slope estimates, we started by assuming that the only knowledge we have is the size ranges of the different groups, but no knowledge of how the biomass of each group is distributed within its size range. To do this, we generated for each group a random allocation of total group biomass across the log-equal bins in its size range, with the total allocation summing to 1. Fig. S12 A and C are the distribution of slopes after 10,000 simulations for the top 200 m, and the entire water column, respectively. For the top 200m, we obtained a normal distribution with mean −0.042 and a standard deviation of 0.006. This gives a 95% confidence interval for the slope of the total global biomass-spectrum in the top 200m of −0.054 to −0.030. For the global biomass-spectrum for the entire water column, we obtained a normal distribution with mean −0.054 and a standard deviation of 0.007. This gives a 95% confidence interval for the slope of average global biomass from bacteria to whales across the entire water column of −0.068 to −0.040.

**Fig. S12.**
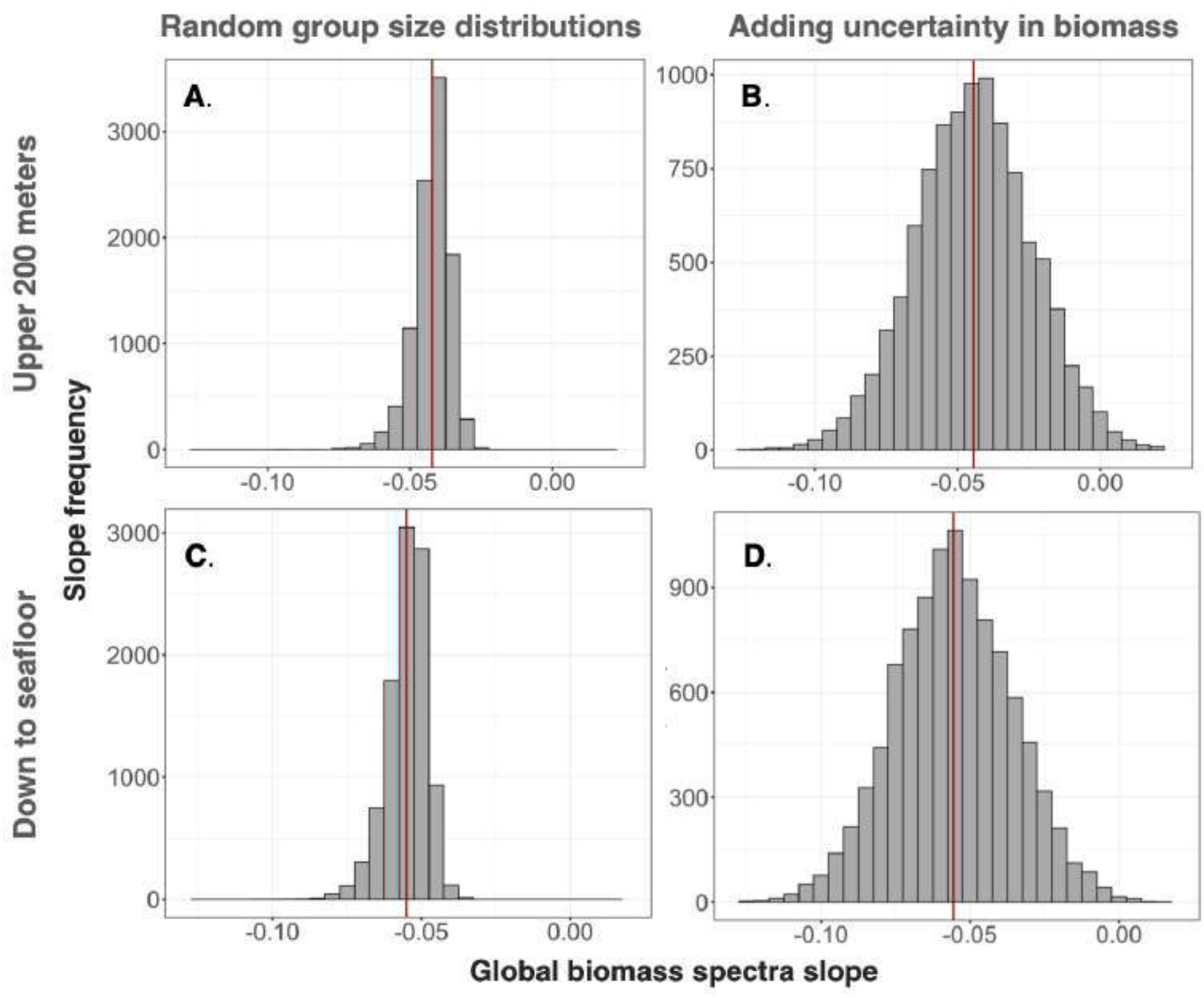
Biomass spectra slopes under different assumptions. Spectra slopes are obtained from 10,000 simulations where we assume no knowledge of the distribution of biomass for each of 12 major groups, listed in Table S2. **A** and **B** show exponent values for the upper 200 meters of the ocean, while **C** and **D** show values for waters to the seafloor. In A and C, we use our best estimates of group biomass, and allocate this at random to different size classes within the group. In B and D, we additionally randomize the assignment of group biomass based on our uncertainty bounds shown in Fig. 2 A.

We not only have uncertainty in the distribution of biomass for each group within its size range, but we also have uncertainty in our estimates of the total biomass for each group. Above, we looked at the uncertainty in the slope of average biomass for each group, but what if we also want to see the additional effect of uncertainty in our estimates of the total biomass for each group? To do this, we added an additional step where for each group we obtained a random total biomass estimate from a log-normal distribution centred at the mean biomass estimate for that group, with standard deviation calculated from the 95% confidence intervals for the group. Fig. S12 B, C are the distribution of slopes for 10,000 simulations for the top 200m and the entire water column, respectively. For the top 200m, we obtained a normal distribution with mean −0.049 and a standard deviation of 0.021. This gives a 95% confidence interval for the slope of the average global biomass in the top 200m from bacteria to whales of −0.085 to −0.003. Across the entire water column, we get a normal distribution with mean −0.056 and a standard deviation of 0.020. This gives a 95% confidence interval for the slope of average global biomass from bacteria to whales across the entire water column of −0.095 to −0.017.

#### Slope dependence on group biomass

Our global maps allow us to estimate the biomass (or abundance) spectrum slope for every grid point over the globe, where all major groups have been estimated. These data exclude mammals, though they represent a small fraction of the total biomass and so may not play so dominant a role in affecting the biomass spectrum. Fig. S13 is a map of the biomass spectra slopes at all the grid points where we have data for each major group from bacteria to fish. We assumed uniform biomass across each major group size range, though as discussed above, this has little effect on the resulting spectra slopes (see previous section). We then summed across major groups in each size class for a total of 33,821 regression slopes. These slopes are almost all in [-0.01, −0.05], which are consistent with the sensitivity analysis described in the previous section (see the histogram of slopes above the map legend in Fig. S13). In other words, the size-spectrum appears to broadly hold globally.

**Fig. S13.**
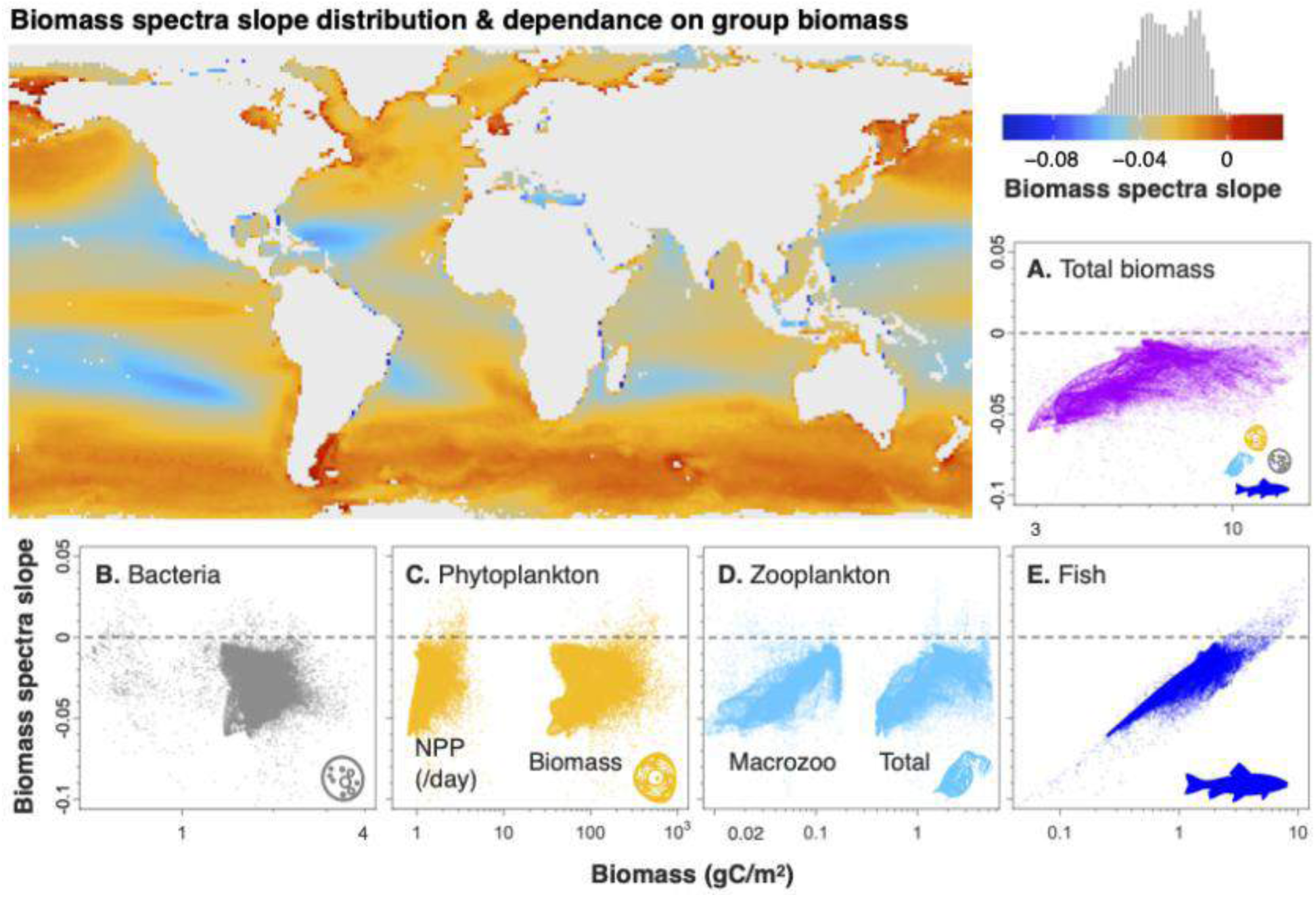
Geographic distribution of biomass spectra slopes and their dependence on major group biomass. The biomass spectra slope is calculated for 33,821 regions in the ocean, with the vast majority of slopes between −0.05 and −0.01, as shown in the legend histogram. The colors in the map denote the slope, while colors in the plots denote major taxonomic groups. For each ocean region, the biomass spectrum slope over all groups is plotted against the biomass of each major group separately, excluding mammals. **A.** Biomass spectrum slope vs. the total biomass sum over all major groups. **B.** Biomass spectrum slope vs. bacteria biomass. **C**. Slope vs. both net primary production (NPP; gC/m^2^/day) and phytoplankton biomass. **D.** Slope vs. macrozooplankton and total zooplankton biomass, the latter of which also includes micro- and mesozooplankton (each of which have similar patterns to the total). **E.** Slope vs. fish biomass. Biomass spectra slopes appear to have a strong, almost proportionate dependence on the logarithm of fish biomass. Fish also exhibit an order of magnitude larger biomass range than other groups.

We also checked how this relatively small variation in slope was related to the biomass of each major group (Fig. S13 B-E). Although there is almost no slope dependence on total biomass, nor on most major groups, there does appear to be a strong slope dependence on fish biomass (Fig. S13 E) in different regions of the ocean. The slope appears to have a strong proportional dependence on the log of fish biomass. Unfortunately, fish also happen to be our least well resolved or constrained group, and so it is difficult to know whether this is representative of biological reality or an artefact of how fish biomass is estimated.

If this slope dependence on fish biomass is legitimate, it indicates the slope gets more negative in more oligotrophic systems, which is consistent with comparative empirical research among plankton (*2*), but contradicts empirical research among fish in northern lakes (*128*). Fig. S13 E implies is that fish get more abundant faster than other groups as we go to more eutrophied systems (and less abundant faster towards oligotrophic). If oligotrophic has a more positive slope than eutrophic (regardless of actual slope values), it follows that fish must have smaller variance than other groups, (regardless of their actual values).

Alternatively, if this slope dependence on fish biomass is an artefact, it implies the fish models are overly sensitive to variations in their prey and are varying more in response than they should in reality. This appears due to fish having the largest range in biomass, spanning more than 2 decades, when other groups span < 1, and that fish occupy the entire upper part of the regression. It is possible that fish biomass exhibits greater variability than other groups over different regions of the ocean due to their aggregation and migratory behaviors, but it is not not clear if that variation would be detected at the coarse grid scale of our global map data. If we could compress this range in biomass variability (without affecting the mean fish biomass) into one order of magnitude, this slope dependence should all but vanish.

### Theoretical considerations

In this section we consider previous research that has attempted to explain the existence of a regular size-spectrum with a slope of −1, including some issues with these attempts. We then proceed to suggest some simple principles that link the size-spectrum with the species level size-density scaling relation, as shown in Fig. 4. Finally we outline some ideas for why the biomass of bacteria and large mammals do not align with other major groups, focusing on energetics.

#### Theoretical background

Since the abundance-size-spectrum slope of −1 was first proposed by Sheldon (*1*), models have proliferated to explain how it might emerge from the myriad species and interactions, life-history strategies, energy pathways, and abiotic and human effects (reviewed in (*2, 5–8*)).

Ideally, a model will be both explanatory and make novel predictions, including those with practical applications, and will be as simple and as general as possible. Despite the many different kinds of models for the size-spectrum, for the most part, there is a core set of underlying processes that are generally assumed in nearly all models. While the size-spectrum is the structural pattern of how biomass is distributed in relation to body mass, all living matter requires energy to be maintained, and is maintained via the dynamics of birth and death, growth and turnover. This means the size-spectrum can be reasoned to be based on either energetic or dynamic processes, and increasingly, is thought to be based on both (*6, 11*).

Most models consider how energy is captured through photosynthesis and transferred to larger bodied individuals through predation, with a given trophic transfer efficiency (the ratio of predator production to that of its prey). Indeed, the predator to prey mass ratio is considered of prime importance in determining the size-spectrum (*5, 6, 8, 22, 23, 129–131*). Further energetic factors such as allometries of metabolism or consumption with body mass are also commonly considered, and assumed to scale near ¾ with body mass. Dynamic processes are also included, either on their own, or in addition to energetics, and consider how individual enlargement from birth to maturity transfers individuals across size classes, with smaller size classes repopulated by reproduction.

Each of these individual-level assumptions are problematic, however, in that they are typically less consistent and general than the size-spectrum itself, the best example being the predator-prey mass ratio. Many size-spectrum models assume a constant predator-prey mass ratio (PPMR; e.g. predators are 500 times more massive than their prey; (*6*). As can be seen from Fig. S14, however, predator-prey mass ratio is not constant, but varies greatly, even for a single individual predator, raising questions about the veracity of this common assumption. Fig. S14 shows three meta-analyses of predator-prey mass ratio, showing that each study shows somewhat different patterns, but that there is no consistent predator-prey mass ratio. Not only are predator-prey mass ratios extremely variable, but there is no consistent slope across studies, much less a constant PPMR (a constant ratio implies a slope of 1). Data for predator-prey mass ratio are from (*21, 22, 132*). Note that Barbier and Loreau 2019 (*132*) included these data as supplemental materials, and the study was not designed to explicitly study predator-prey mass relation.

**Fig. S14.**
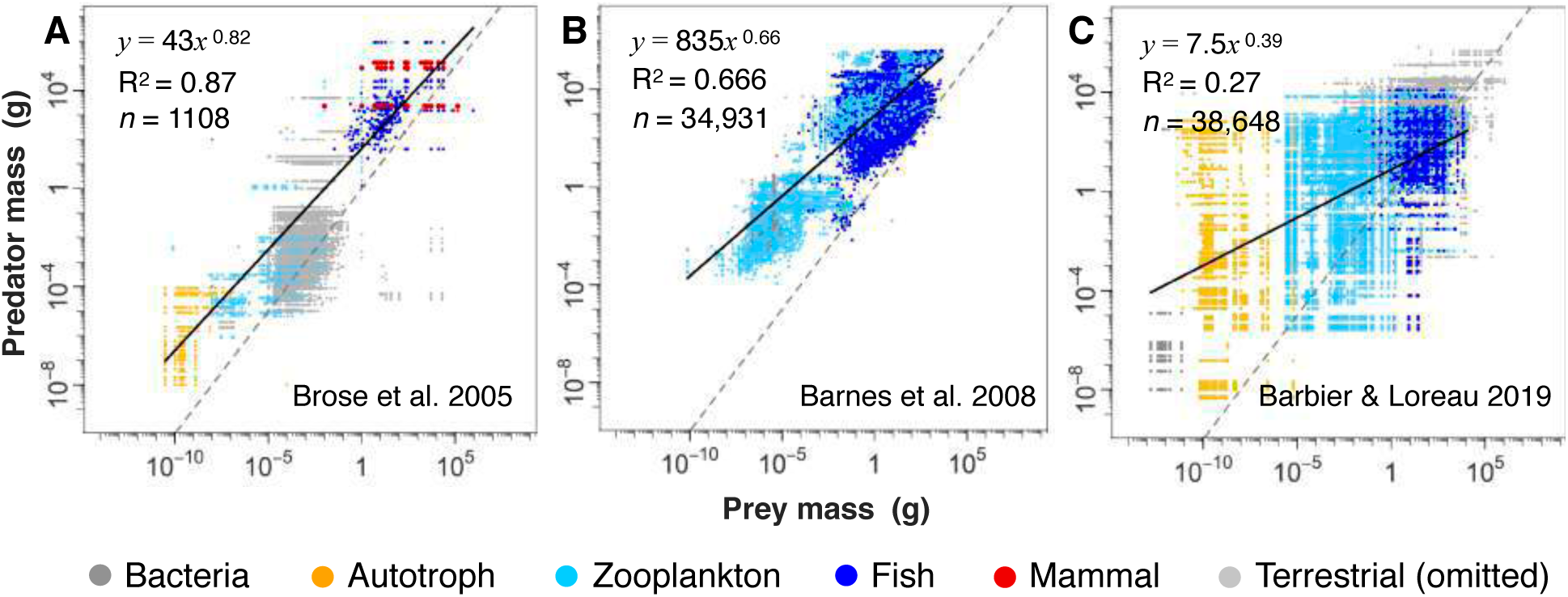
Predator-prey mass relations. In all three plots, the typical mass (grams) of different species consumer-resource pairs are plotted. Only aquatic trophic interactions are colored (according to prey), for which the best fit regression (black line) equation is reported. The dashed grey line is the 1:1 line, meaning a predator consumes a prey of the same size, with points below the dashed line representing a smaller individual consuming a larger. Data are from **A**. Brose et al. 2005, **B.** Barnes et al. 2008, and **C.** Barbier & Loreau 2019. The level of overlap between studies has not been investigated, but the patterns are all very highly dispersed.

Other energetic assumptions include the rates of individual metabolism and consumption. A number of proposed explanations for the size-spectrum assume a constant predator-prey mass ratio and trophic transfer efficiency, a constant population-level energy use (termed energetic equivalence and discussed in the sub-section below), and near ¾ metabolic scaling with body mass (*133–136*). In addition to the problems posed by the assumption of a constant predator-prey mass ratio, and further challenges raised below to the assumption of energy equivalence, metabolism does not scale near ¾ across the size-spectrum, but scales closer to linear (exponent of 1; (*10, 24*)). Even within major groups like phytoplankton, zooplankton or fish, metabolic scaling is considerably elevated above ¾ (*10*).

In addition to energetic factors thought to underly the size-spectrum are dynamic processes that move individuals across size classes through growth, turnover, reproduction and mortality (*5, 137*). These dynamics are particularly pronounced in fish, which produce extreme numbers of tiny offspring, most of which die early in ontogeny, but some of which pass through a great many size classes through their life history (*25, 138*). This life-history contrasts markedly with mammals and even many crustaceans that produce a relatively small number of offspring that are a constant fraction of the adult size (about 1/100; (*25*)). These life-history differences between groups imply that a single general size-spectrum theory would be unlikely to be based on the unique dynamics of each group.

While these individual-based processes are undoubtedly related to the size-spectrum, as has long been assumed, the order of causality is less obvious. While most modeling studies assume that these individual level relations determine the community-level size-spectrum, the assumptions do not appear to be as robust as was once thought. The predator-prey mass ratio is not a constant ratio, but is extremely variable (Fig. S14), the scaling of metabolism and consumption is not actually what is widely assumed (slope near ¾), but significantly steeper (*10, 24*), and many of the life-history traits that may be thought to move biomass across size classes, tend to be group specific, with fish exemplifying unique life-history traits. Indeed, most of the commonly assumed underlying processes and body mass allometries for the size-spectrum are neither as general or as consistent as the size-spectrum itself and thus fail to account for its overall regularity from bacteria to whales.

From examination of the body mass allometry data commonly used to posit an underlying basis for the size-spectrum, we find that few of these allometries exhibit the patterns that are so often claimed by size-spectrum theory (reviews in (*2, 5–8*)). We argue that current theory has not identified the fundamental causal drivers, and as such is unable to fully explain the full generality and apparent regularity of the size-spectrum.

Two broad exceptions that may provide further insight into the size-spectrum are bacteria and mammals, both of which appear to defy the overall biomass constancy. As shown in Fig. 2, bacteria and mammals appear to be exceptions to the general constancy in biomass exhibited by other groups. These exceptions become more pronounced when we consider the size-spectrum down to the seafloor (further relative increase in bacteria; Fig. 2) or if we consider the modern-day size-spectrum, that includes two centuries of cumulative human impacts (further relative decrease in mammals; Fig. 4). In the sections that follow we examine possible explanations for these exceptions. We first consider metabolic requirements of major groups, and finally describe human impacts on the size-spectrum, as two possible avenues that may account for these exceptions.

#### Links to size-density scaling

A closely related pattern to the size-spectrum is the species-level size-density relation (also known as abundance-mass scaling; (*9, 10, 89, 94, 95, 139, 140*). This is the population density of a species in its natural habitat within its geographic range area plotted against mature body mass. For several decades abundance density was thought to scale with an exponent near −¾ to body mass, and within particular taxonomic groups such as mammals, this appears to hold, but across all groups, it was recently shown that the exponent is closer to −1, mirroring in inverse fashion the metabolic exponent both within and across major groups (*10*).

As we show in Fig. 4, after re-scaling the size-spectrum for the global ocean (Fig. 1) by total ocean area, we can express the size-spectrum in terms of size class density, which allows the size-spectrum to be directly compared on the same axes as the data for size-density scaling. Data for the size-density relation are from (*10*) (with terrestrial species removed), and are updated with data for marine mammal species obtained by this study. Other than these marine mammal data, the data sources for the size-spectrum and size density scaling shown in Fig. 4 are entirely independent. The fact that they so closely parallel one another, with an approximately order of magnitude offset in elevation, is thus surprising.

Despite the similarities shown in Fig. 4, however, there are some important differences between size-spectra and size-density scaling. The size spectra refers to a single community such as a lake or region of the ocean, while the size-density relation encompasses very different aquatic communities. Moreover, the size spectra is the sum of all abundances of all species within a size class, while the size-density relation is concerned with the abundance of a single species of a given adult size within its geographic range area. The average adult size and abundance of a species of fish is also quite different from the actual sizes and abundances of larvae, juveniles, and adults of that same species residing in a community, and though large fish are comparatively very rare, they produce a very large number of small offspring (*138*), most of which are consumed as juveniles (Fig. S15). Some communities may also not have equal diversity across logarithmic size classes. The connections between the within-community size-spectra and the cross-community size-density relation is thus less straightforward than Fig. 4 might suggest, with both relations being constructed in very different ways from independent sources.

**Fig S15.**
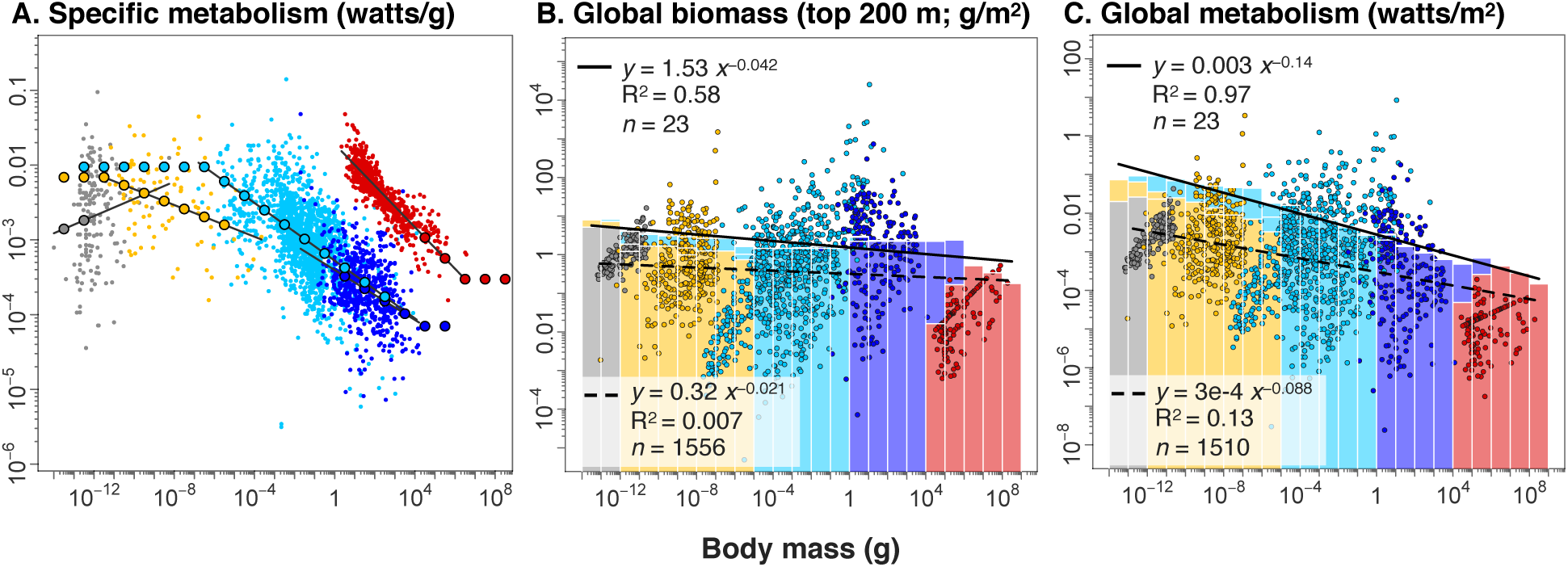
Ocean energetics. **A.** Mass-specific metabolism (watts/g) are plotted against body mass, and major group level regressions are used to estimate specific metabolism for each group in each size-class (large circles). Where estimates are needed beyond the range of the data available, we have used the closest regression value within the data range, rather than extending the regression beyond the data range. **B.** The global biomass density spectra for the top 200 meters of ocean are compared against species level biomass density (g/m^2^) from data in Fig. 4. Slightly negative slopes are apparent due to higher than average bacteria biomass and lower than average marine mammals. **C**. The global energy density spectrum is compared against species-level population energy use (both in watts/m^2^). Both relations are obtained by multiplying the respective biomass densities in B with metabolism per gram in A, and show more negative slopes than in B.

One way to connect these patterns is that the size-spectrum (frequency of individuals in a size class) is equal to the product of i) the mean species density (per size class), ii) the species diversity (per size class), and iii) the mean fraction of total area occupied by species geographic ranges (per size class). The concordance between the size-spectrum and size-density scaling suggests that diversity and geographic range area may scale with size, or, or else trade-off against one another.

#### The ocean energy spectrum

The observation that bacteria are above and marine mammals below the otherwise constant biomass spectrum of all other taxonomic groups could indicate that energetics may be playing an important role in the shape of the biomass spectrum. Across eukaryotes, representing more than 20 orders of magnitude in body mass, species population metabolism is known to vary within only about three orders of magnitude, while population biomass is considerably more variable (*10*). This is generally known as the ‘energetic equivalence’ rule and is calculated by multiplying species population density by their basal metabolism, which both scale with body mass with exponents that are inverse to one another (*9, 10, 89, 94, 95, 139, 140*).

Bacteria are well known to enter severely reduced energy states by taking highly resistant and inactive forms (*141*). These states have almost negligible rates of respiration and may represent a dominant state of most bacteria in the deep ocean, consistent with their very slow turnover times (*31*). On the other hand, endothermic mammals have 1-2 orders of magnitude higher metabolic rates than ectotherms of the same body mass (*10, 24*). We thus reasoned that the higher than expected bacteria biomass (Fig. 2), multiplied by their overall lower metabolic states, and the lower than expected mammal biomass multiplied by their higher metabolic rates, might better align these two groups with the energetics of all other taxonomic groups.

To test this hypothesis, we assembled species-level mass-specific metabolism (from (*10, 11*)) and fit body mass regressions to all major taxonomic groups. For each order of magnitude size-class, and each taxonomic group, we estimated a mass-specific metabolic rate (large circles in Fig. S15 A). For size-classes that extended beyond the size range of our regressions, we used the closest value of metabolism which the regression predicted, rather than extrapolating the regression beyond its known size range. These mass-specific metabolism values were then multiplied by the biomass for each size class in the upper 200 meters of the ocean (Fig. S15 B). Also shown in Fig. S15 B are species-level estimates of biomass for aquatic species (see Fig. 4; data from (*10*)). The resulting energetic spectrum is shown in Fig. S15 C and reveals a significantly decreasing size-class metabolic rate with increasing size, with a slope of −0.14. Although this does indeed align better the energetic demands of bacteria and mammals with the other taxonomic groups, the significant downward slope is not consistent with an energetic equivalence across the size-spectrum. It is somewhat surprising that marine species level population metabolism (points in Fig. S15 C) also reveal a shallow negative slope with body mass, contrasting with the energetic equivalence seen among all species (both terrestrial and aquatic) combined (*10*).

Our findings suggest that the size-spectrum does not appear to be fundamentally underpinned by energetic considerations, as the total energy use per size class actually destroys the invariance: marine metabolism has a slope near −0.14. Despite better aligning bacteria and mammals with all other groups, they are aligned along a significantly diminishing relation, rather than presenting a more robust invariance. This contrasts with our hypothesis that the unique metabolic states of bacteria and mammals may compensate their relative offsets in biomass. Given that common energetic assumptions about metabolic and consumption scaling (assumed to be near ¾, but are actually closer to 1) and the predator-prey mass ratio (assumed constant, but actually highly variable) are so often mistaken, we take these findings to indicate that any strictly individual energetic basis to the size-spectrum needs to be re-evaluated.

In this section, we have considered the empirical basis for many size-spectrum theories, showing that many common assumptions, particularly with respect to energetics are not well founded, or do not apply across the size range. Finally, we have considered how bacteria and mammals offer exceptions to the biomass invariance that may offer insight into the origin of this otherwise regular and consistent pattern, but the most obvious energetic factors do not elucidate the basis for the size-spectrum. This leaves the question unsolved, but hopefully better delineates the contours of the problem.

